# Imipramine and olanzapine block apoE4-catalyzed polymerization of Aβ and show evidence of improving Alzheimer’s disease cognition

**DOI:** 10.1101/2021.03.28.437389

**Authors:** Noah R. Johnson, Athena C.-J. Wang, Christina Coughlan, Stefan Sillau, Esteban Lucero, Lisa Viltz, Neil Markham, Cody Allen, A. Ranjitha Dhanasekaran, Heidi J. Chial, Huntington Potter

**Affiliations:** University of Colorado Alzheimer’s and Cognition Center, Linda Crnic Institute for Down Syndrome, Department of Neurology, University of Colorado, Anschutz Medical Campus, Aurora, CO, USA

## Abstract

The apolipoprotein E (APOE) ε4 allele confers the strongest risk for late-onset Alzheimer’s disease (AD) besides age itself, but the mechanism(s) underlying this risk are debated. The critical test of any proposed AD mechanism is whether it leads to effective treatments. We developed a high-throughput assay to identify inhibitors of apoE4-catalyzed polymerization of the amyloid β (Aβ) peptide into neurotoxic fibrils. Screening a human drug library, we identified five non-toxic, blood-brain-barrier-permeable hit compounds that reduced apoE4-promoted Aβ and tau neuropathology in cultured neurons. Two hit compounds, imipramine and olanzapine, but not other (non-hit) antipsychotics or antidepressants, when prescribed to AD patients for their normal clinical indications, led to improvements in cognition and clinical diagnosis. Imipramine and olanzapine have no structural, functional, or clinical similarities other than their ability to inhibit apoE4-catalyzed Aβ polymerization, thus identifying this mechanism as an essential contribution of apoE4 to AD.

**One Sentence Summary:** High-throughput drug screens, studies in Alzheimer’s disease cell culture models, and analyses of human clinical data identified inhibitors of the apoE4-Aβ interaction as a novel class of Alzheimer’s disease therapeutics.

## Introduction

Alzheimer’s disease (AD) is the most common neurodegenerative disorder, currently afflicting 5.8 million Americans and 50 million people globally, and these numbers are projected to grow to 13.8 million and 152 million, respectively, by 2050 (*1*). The amyloid cascade hypothesis is supported by genetic, biochemical, and pathological data and posits that AD is initiated by oligomers of the amyloid-β (Aβ) peptide, which is formed by sequential cleavage of the amyloid precursor protein (APP) by β- and γ-secretases (*2*). Increased production and/or impaired clearance of Aβ leads to its accumulation as amyloid plaques in the brain parenchyma and vasculature, followed by intraneuronal aggregation of the microtubule-associated protein tau into neurofibrillary tangles (NFTs) and neuronal cell death (*3*). Accordingly, numerous therapeutics targeting the amyloid cascade have been developed, such as secretase inhibitors and immunotherapies directed against Aβ or tau (*4*). However, AD therapies have a very poor track record in human clinical trials. As a result, none of the currently approved AD treatments can reverse, or even substantially slow, the progression of disease, underscoring the importance of alternative mechanistic approaches to treatment.

Genetic factors can increase the risk for developing AD, in particular in individuals who carry the ε4 allele of the apolipoprotein E (*APOE*) gene (*5*). The three common apoE isoforms, apoE2, apoE3, and apoE4, differ by single amino acid substitutions at positions 112 and 158. Of the three common allelic variants of *APOE*, ε3 is most prevalent, accounting for 70-80% of the total alleles in the human population, followed by ε4, which accounts for 10-15%, and then ε2, which accounts for 5-10% (*6*). Carrying one copy of *APOE4* more than triples the risk for AD, whereas being homozygous for *APOE4* increases the risk by greater than 12-fold (*5*). Indeed, despite its low allelic frequency in the general population, approximately 60-65% of individuals with AD carry at least one copy of *APOE4* (*7*). The onset of AD symptoms occurs earlier in *APOE4* carriers than in non-carriers and is accompanied by more severe plaque deposition, intraneuronal Aβ accumulation, cerebral amyloid angiopathy, and blood-brain barrier (BBB) dysfunction (*5, 8, 9*).

Human apoE is a 34 kDa glycoprotein that has pleiotropic functions including in both the central nervous system (CNS) and the peripheral nervous system. In the CNS, apoE is primarily expressed by astrocytes and microglia and serves as the primary carrier of cholesterol and other lipids (*10*). ApoE also functions as a modulator of synaptic plasticity, neuroinflammation, glucose metabolism, and vascular integrity in the brain (*11*). The expression of apoE is upregulated in the AD brain, particularly in regions where amyloid deposits first, concurrent with an inflammatory cascade caused by toxic Aβ oligomers [reviewed in (*12–14*)]. Multiple potential mechanisms by which apoE4 increases the risk for AD have been proposed and investigated. For example, apoE, and especially apoE4, binds to Aβ with high affinity and acts as a catalyst to accelerate the rate of Aβ oligomer and fibril formation (*15–18*), increase their stability (*19, 20*), and promote their neurotoxicity (*21–23*). Consistent with this premise, human apoE4 expressed in mice seeded Aβ aggregation (*24*), and conversely, knockout (KO) of the mouse *Apoe* gene in transgenic mice expressing human APP abolished amyloid fibril and plaque formation and cognitive decline (*25, 26*). Furthermore, careful longitudinal evaluation in prodromal AD has revealed that *APOE* genotype plays the greatest role during the initial seeding stages of Aβ deposition (*27*). Additional contributors to the increased genetic risk of *APOE4* in AD may include impaired Aβ clearance, exacerbated oxidative stress and neuroinflammation [reviewed in (*28–30*)], and loss of critical apoE functions. Notably, apoE is found co-deposited in amyloid plaques in the AD brain, suggesting a direct interaction with Aβ (*31*). Rare mutations in the Aβ binding domain of apoE markedly reduce the risk for AD in humans (*32, 33*). Taken together, substantial evidence supports a role for apoE as an essential molecular chaperone for Aβ aggregation in the brain, and suggests that inhibiting this process is a promising therapeutic approach to preventing AD.

ApoE-targeted therapeutics for AD have focused predominantly on modulating the overall levels of apoE or the degree of its lipidation. Therapeutic apoE depletion has been evaluated in mice using antisense oligonucleotides or immunotherapies to reduce apoE levels or inhibit its function (*34, 35*). Given that apoE is expressed throughout the body where it carries out many critical functions, a reduction in total apoE levels is expected to have many undesirable side effects (*36*). Thus, focusing on the interaction between apoE and Aβ may yield a more precise therapeutic benefit for AD without interfering with the many beneficial functions of apoE. Small molecule ‘structure correctors’ or gene editing have been used to block the formation of the pathological conformation of apoE4 (*37, 38*). Additionally, a synthetic peptide corresponding to the apoE-binding site on Aβ was also found to reduce Aβ aggregation *in vitro* and in AD mouse models (*21, 39*). Although the clinical translatability of these therapies remains to be determined, together they validate the inhibition of the apoE4-Aβ interaction as a tractable therapeutic approach for AD.

Here, we describe the identification of a set of small molecule drugs that can block the interaction between apoE4 and Aβ. We developed an apoE4-catalyzed Aβ fibrillization assay and employed it for high-throughput screening (HTS) of the National Institutes of Health (NIH) Clinical Collection (NCC) library of small molecules with a history of use in clinical trials. Repurposing known drugs has numerous benefits, such as the availability of safety and dosing information that allows for faster and cheaper clinical testing. Through a series of HTS assays, we identified eight hit compounds that reduced apoE4-catalyzed Aβ fibrillization in a dose-dependent manner. We present evidence that two of those hit compounds− imipramine and olanzapine− reduced Aβ and phosphorylated tau (pTau) neuropathology in cell culture models, and when taken by AD patients for their other normal clinical indications, were associated with improved cognition and greater incidence of receiving an improved clinical diagnosis. Because imipramine and olanzapine are completely different from each other with regard to their structures, designed mechanisms of action, and current approved clinical indications, and their only common feature is our discovery of their shared ability to block the apoE4-catalyzed polymerization of Aβ into neurotoxic fibrils, these findings validate this mechanism as an essential contribution of apoE4 to AD.

## Results

### Development of an apoE4-catalyzed Aβ fibrillization assay for HTS

Building on earlier work (*15, 16, 19, 21*), we adapted an Aβ fibrillization assay monitored with the amyloid-binding dye thioflavin T (ThT) to study the catalytic effects of apoE4 and optimized it for HTS for inhibitors of the apoE4-Aβ interaction. Utilizing a design of experiments (DOE) approach, we first determined the optimal concentrations of Aβ42, apoE4, and ThT to generate a dose-responsive readout. In our initial experiment, we found that lowering Aβ concentration resulted in reduced Aβ fibrillization rate and growth phase duration (Fig. 1A and Fig. S1). We confirmed this result in a second experiment, and also found that the baseline level of ThT fluorescence could be reduced by decreasing the concentration of ThT; however, a higher concentration was necessary to observe the maximal ThT fluorescence readout (Fig. 1B and Fig. S2). We also observed that 1 nM apoE4 resulted in greater Aβ fibrillization than did higher concentrations of apoE4, although the effect could be overcome by increasing the concentration of Aβ (Fig. S3), suggesting that the Aβ:apoE4 ratio was important. In a third experiment, we found that indeed, 1 nM apoE4 accelerated Aβ fibrillization but that increasing apoE4 to 2 nM negated its catalytic effect (Fig. 1C). Consistent with these findings, the integrated area under the curve (AUC) of ThT fluorescence increased with greater quantity of Aβ used (Fig. 1D), while the fold-change in ThT fluorescence increased with a greater Aβ:apoE ratio (Fig. 1E). Finally, a response optimization algorithm was used to identify the concentrations that maximized both the AUC and the fold-change of ThT fluorescence simultaneously, which was determined to be 20.9 μM Aβ42, 0.75 nM apoE4, and 14.8 μM ThT (Fig. 1F and Fig. S4).

**Fig. 1.**
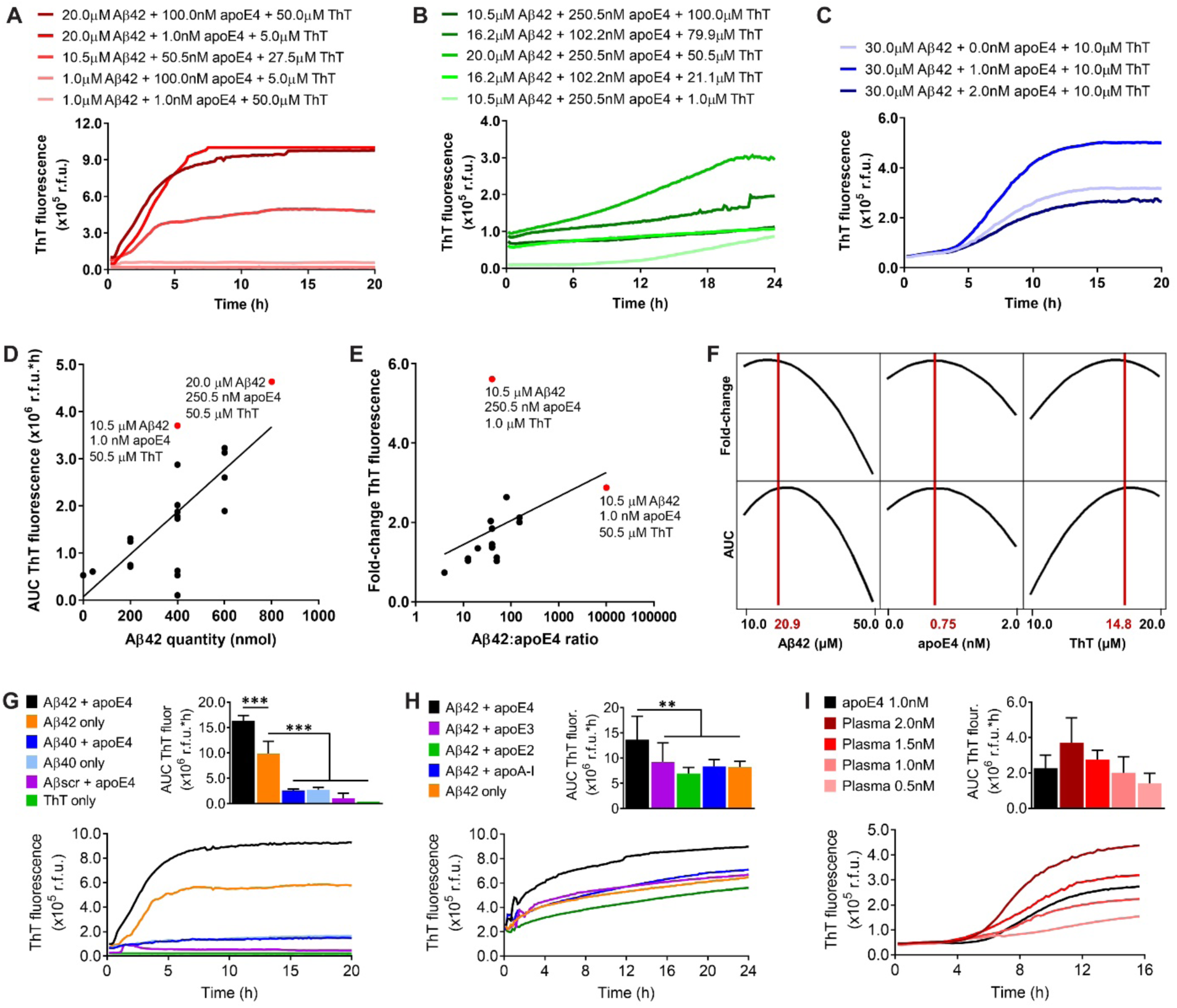
Development of an apoE4-Aβ fibrillization assay for HTS. (**A**) Concentrations of Aβ42, apoE4, and ThT were varied in a ½ fraction factorial experiment in a 96-well plate and ThT was measured in relative fluorescence units (r.f.u.) in n = 3 wells per group. (**B**) Concentrations of Aβ42, apoE4, and ThT were varied in a response surface experiment in a 384-well plate in n = 3−4 wells per group. (**C**) Concentrations of Aβ42, apoE4, and ThT were varied in a second response surface experiment in a 384-well plate in n = 6 wells per group. The experiment was replicated twice and the results were combined. (**A**−**C**) Several representative groups were plotted and the complete results are provided in Data file S1. (**D**) The quantity of Aβ42 was plotted against the area under the curve (AUC) of ThT fluorescence in r.f.u.*hours (h). Linear regression was performed to identify a best-fit line (R^2^ = 0.53). Two assay conditions that maximized the AUC were identified (red dots). (**E**) The Aβ42:apoE4 ratio was plotted on a log scale against the fold-change in ThT fluorescence. Linear regression was performed to identify a best-fit line (R^2^ = 0.14). Two assay conditions that maximized the fold-change were identified (red dots). (**F**) In the second response surface experiment, the optimal concentrations of Aβ42, apoE4, and ThT that maximize both the AUC (r.f.u.*h) and the fold-change in ThT fluorescence were identified (red lines). (**G**) The specificity of the optimized apoE4-Aβ fibrillization assay was evaluated for Aβ42, Aβ40, and a scrambled Aβ42 peptide (Aβ_scr_), with and without apoE4, or for ThT only. The data represent the mean ± SD of n = 3 wells per group. \*\*\**P* < 0.001 by one-way ANOVA. (**H**) The effect of each human apoE isoform, or apoA-I, at 1 nM concentration on Aβ42 fibrillization were evaluated. The experiment was replicated twice and the results were combined. The data represent the mean ± SD of n = 7 wells per group. \*\**P* < 0.01 by one-way ANOVA. (**I**) The effect of apoE isolated from human plasma at different concentrations on Aβ42 fibrillization was compared with that of recombinant apoE4. The data represent the mean ± SD of n = 4 wells per group.

We next evaluated the specificity of our optimized Aβ fibrillization assay. We found that replacing Aβ42 with Aβ40, which is two amino acids shorter, or using a scrambled sequence consisting of the same 42 amino acids (Aβscr), each resulted in significantly less fibrillization (Fig. 1G), consistent with prior reports (*40*). We also tested the effect of other apolipoproteins and found that only apoE4 catalyzed Aβ42 fibrillization, while apoE3, apoE2, and apolipoprotein A-I, did not (Fig. 1H). To confirm that our results with recombinant human apoE4 were translatable to a normal human population, we then tested apoE isolated from pooled human plasma that contained a mixture of all three apoE isoforms. We found that human plasma-derived apoE catalyzed Aβ42 fibrillization in a dose-dependent manner and at a similar level to that of recombinant apoE4 (Fig. 1I). Finally, to verify the usefulness of our assay for HTS of drug libraries, we added different concentrations of DMSO and found no significant effect up to 10% v/v (Fig. S5).

### HTS identifies small molecule inhibitors of apoE4-catalyzed Aβ fibrillization

The NCC drug library contains small molecule compounds that have a history of use in human clinical trials. We performed an exploratory drug screen of 595 compounds from the NCC library, testing each compound at a concentration of 2 µM, and we identified 134 hits (Fig. S6 and Data file S2). We then performed a literature search to determine whether the hit compounds or their metabolites had been reported to be capable of crossing the BBB. Of the 134 hits, we found credible reports that 87 of the compounds had good BBB permeability (Data file S2). We next analyzed the dose-response effects of these 87 compounds on the kinetics of apoE4-catalyzed Aβ fibrillization in our optimized HTS assay. We identified eight hit compounds using the criteria that they reduced apoE4-catalyzed Aβ fibrillization by at least 30% and generally displayed a dose-dependent effect (Fig. 2A). The eight hit compounds ranged in size from 214−580 Da and had varied chemical structure, although every compound contained at least one aromatic ring (Table 1). The hit compounds appeared to reduce the fibrillization rate, resulting in a lower quantity of Aβ fibrils after 24 h (Table 1). Notably, sulfacetamide, EGCG, PD 81723, and indirubin had effect at sub-micromolar concentrations. EGCG was previously shown to inhibit Aβ aggregation in rodent models of AD (*41*) and is currently being tested in human clinical trials for AD (e.g., NCT03978052), thus validating our overall screening approach.

**Fig. 2.**
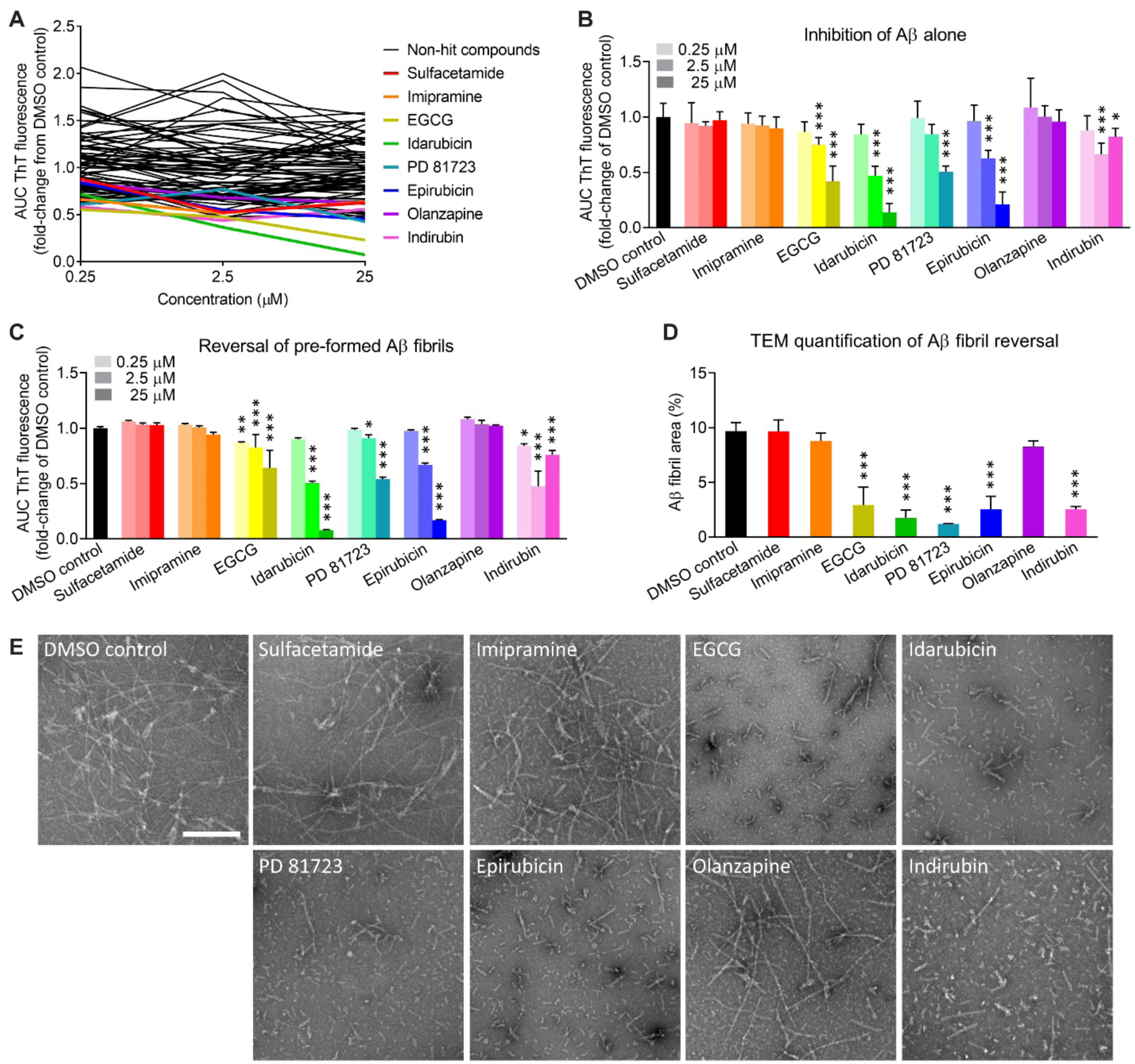
Identification of small molecule compounds that inhibit the apoE4-Aβ interaction or reverse Aβ42 fibril formation. (**A**) HTS of 87 compounds was performed. Small molecule compounds were added at 0.25, 2.5, and 25 µM in a final concentration of 5% (v/v) DMSO. Eight hit compounds (colored lines) were identified. The data represent the mean of n = 3 wells per concentration for each compound, relative to the mean of n = 8 wells for the DMSO control. (**B**) Eight hit compounds were tested for inhibition of Aβ42 fibrillization independent of apoE4. (**C**) Eight hit compounds were tested for disaggregation of pre-formed apoE4-catalyzed Aβ42 fibrils. (**B** and **C**) The experiment was replicated twice and the results were combined. The data represent the mean ± SD of n = 8 wells per concentration for each compound. \**P* < 0.05, \*\**P* < 0.01, and \*\*\**P* < 0.001 compared to the DMSO control by one-way ANOVA. (**D**) ApoE4-catalyzed Aβ42 fibrils were treated for 30 min with DMSO or with each compound at 25 μM, except for compound 077, which was tested at 2.5 μM because it had the greatest effect in previous experiments. TEM images were acquired and analyzed for the Aβ fibril area (%). The data represent the mean ± SD of n = 3 separate TEM images for each compound. \*\*\**P* < 0.001 compared to the DMSO control by one-way ANOVA. (**E**) Representative TEM images of apoE4-catalyzed Aβ42 fibrils treated with DMSO or with each hit compound. Aβ fibrils are observable by negative-stain TEM as light objects on a dark background. Scale bar = 200 nm.

**Table 1.**
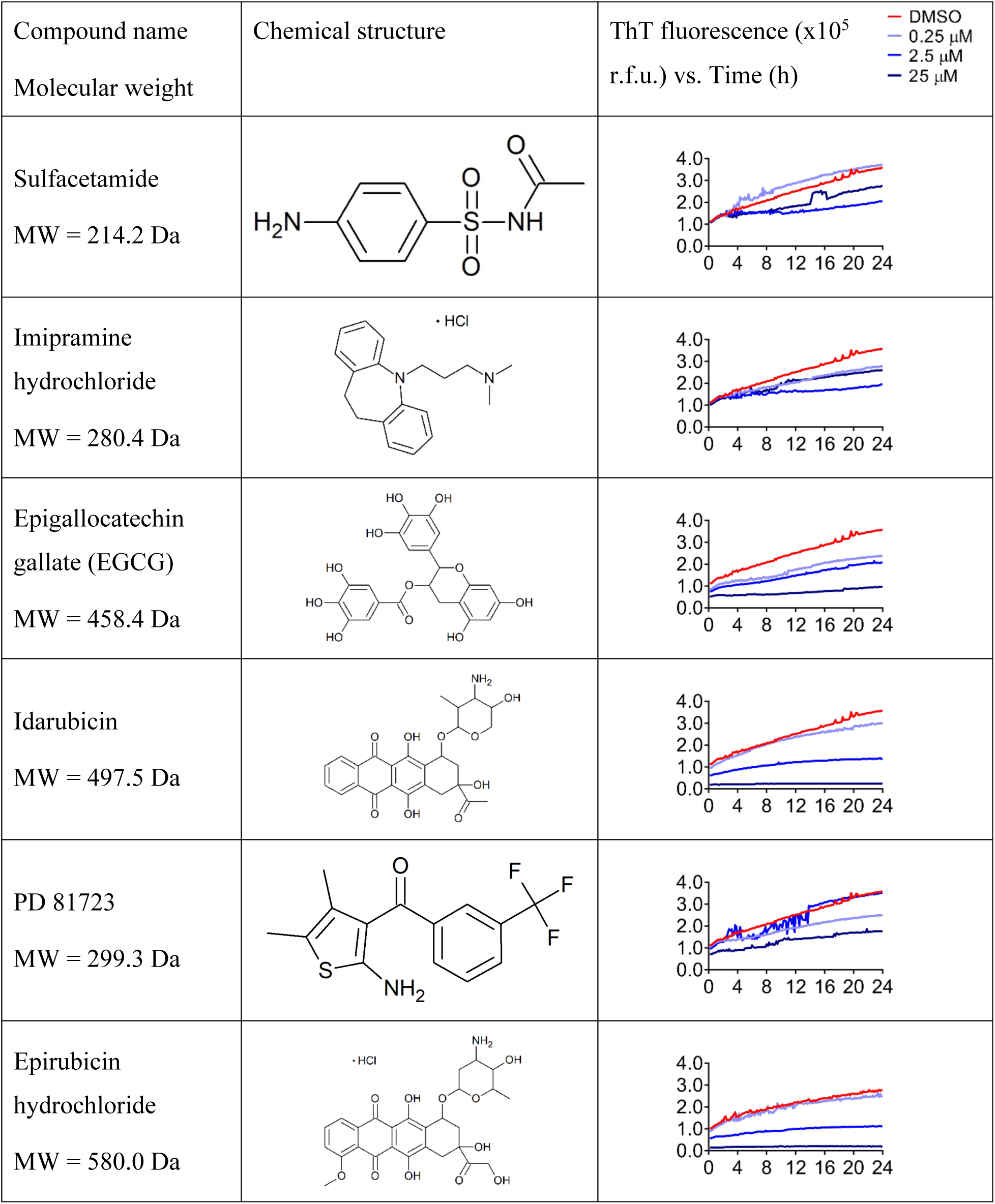
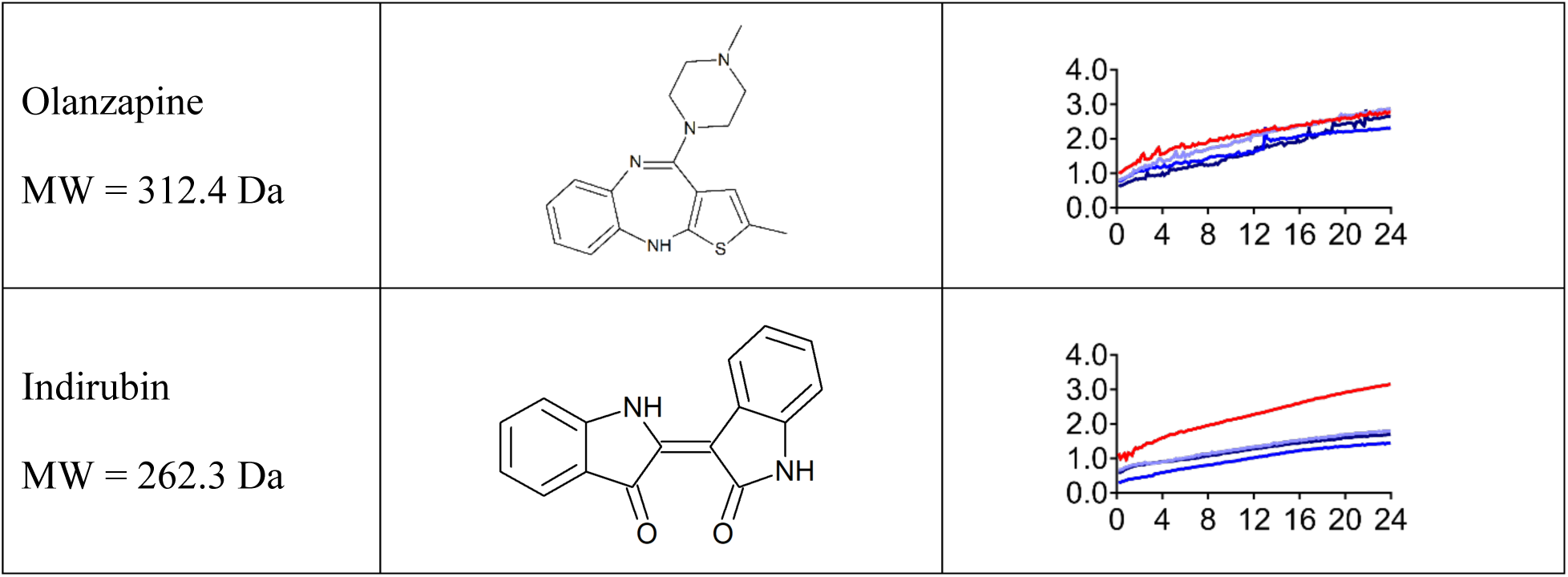
Small molecule inhibitors of apoE4-catalyzed Aβ42 fibrillization.

We sought to identify which compounds were blocking the apoE4-Aβ interaction and which compounds were acting directly on Aβ. EGCG, idarubicin, PD 81723, epirubicin, and indirubin inhibited the fibrillization of Aβ42 alone, independent of apoE4 and in a largely dose-dependent manner (Fig. 2B), suggesting that these five compounds act directly on Aβ. In contrast, sulfacetamide, imipramine, and olanzapine had no effect on the fibrillization of Aβ alone, suggesting that these three compounds are specific inhibitors of the apoE4-Aβ interaction. We also tested the ability of all eight compounds to reverse Aβ fibrillization by first pre-forming apoE4-catalyzed Aβ fibrils and then treating them with each compound. We found that only the five compounds that acted directly on Aβ (i.e., EGCG, idarubicin, PD 81723, epirubicin, and indirubin) could reverse Aβ fibrillization (Fig. 2C). Finally, we used transmission electron microscopy (TEM) to confirm that these compounds disaggregated Aβ fibrils (Fig. 2D), rather than preventing ThT binding or fluorescence. In contrast to the numerous, long Aβ fibrils present following treatment with DMSO, we observed much shorter and fewer Aβ fibrils and aggregates following treatment with the reversal compounds (Fig. 2E). These molecules may be pursued as interventional treatments for patients with pre-existing AD neuropathology.

### Small molecule compounds reduce Aβ neuropathology in primary neurons from 5xFAD transgenic mice

We used an *in vitro* primary neuron assay to examine the cytotoxicity of the small molecule hit compounds as well as their efficacy at reducing intracellular and extracellular Aβ neuropathology under conditions more closely resembling the physiological concentrations of Aβ and apoE than were used in HTS (**Fig. 3A**). Primary neurons isolated from 5xFAD transgenic mice, which express human APP with three familial AD mutations and also express the human presenilin 1 gene (PSEN1) with two familial AD mutations (*42*), were exposed to Aβ42 and/or apoE4, or phosphate buffered saline (PBS) as a negative control. Aβ neuropathology developed in the form of intracellular and extracellular Aβ aggregates in cells exposed to Aβ alone or exposed to apoE4+Aβ, whereas no Aβ neuropathology was observed in cells exposed to apoE4 alone or to PBS (**Fig. 3B**). It is worth noting that human APP and PSEN1 expression in transgenic mouse neurons begins prior to birth, and that the Aβ neuropathology observed is likely comprised of both the Aβ added as a seed and the Aβ produced by the cells. Quantification of cell nuclei revealed that exposure to apoE4+Aβ resulted in a significant reduction in cell viability compared to PBS, apoE4 alone, or Aβ alone (**Fig. 3C**). Significantly more Aβ neuropathology was also present in cells exposed to apoE4+Aβ compared to Aβ alone (**Fig. 3D**), suggesting that apoE4 catalyzes Aβ fibril formation in cell culture medium as it does in acellular assays and that the resulting Aβ fibrils are neurotoxic.

**Fig. 3.**
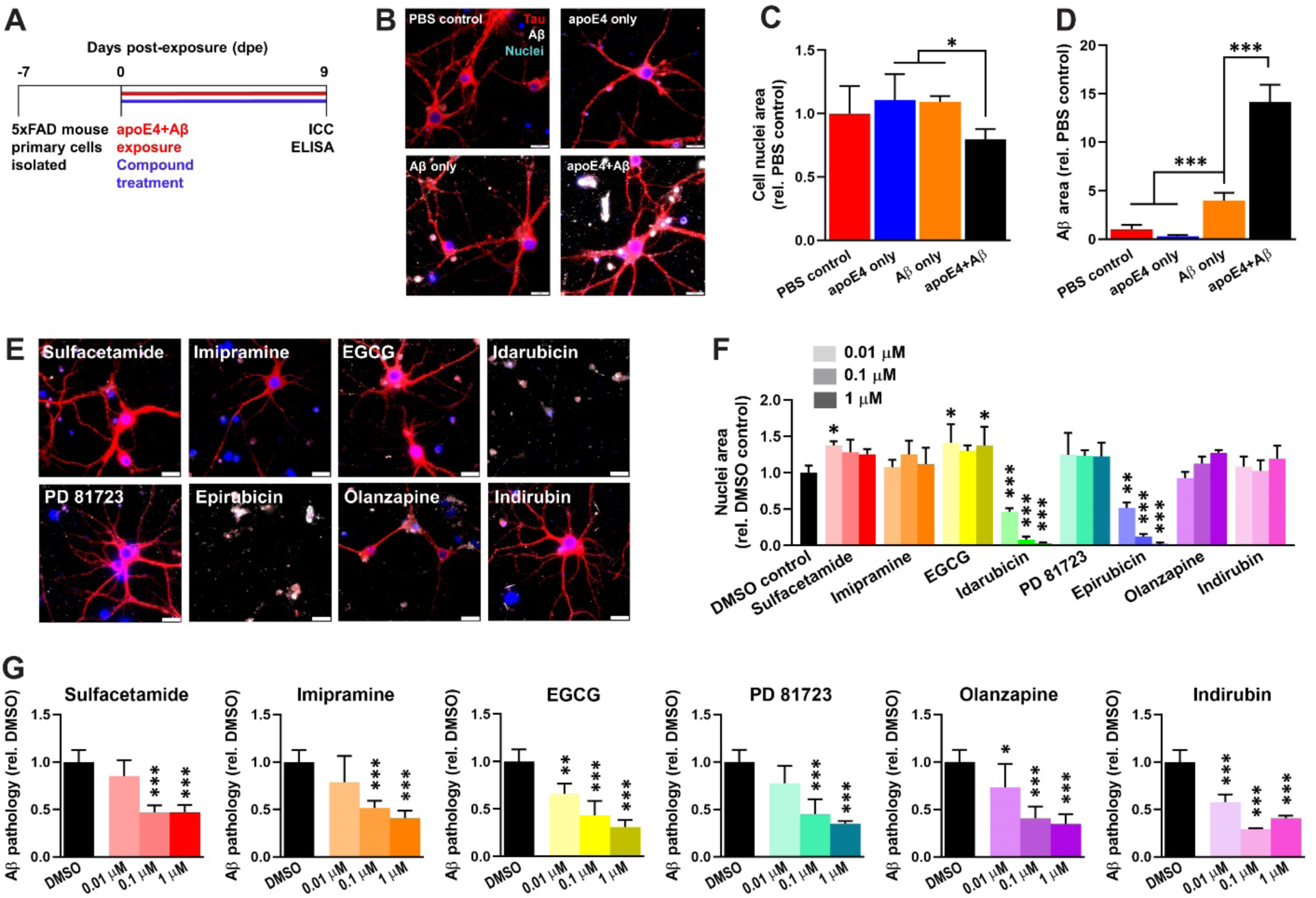
Small molecule compounds inhibit apoE4-catalyzed Aβ pathology in primary neurons from 5xFAD transgenic mice. (**A**) Schematic of drug efficacy experiments using primary neurons from the 5xFAD transgenic mouse model of Alzheimer’s disease. One week after cell isolation from day P1-P2 pups, cells were exposed to 100 nM Aβ42 and 1 nM apoE4 and were treated concurrently with 0.01, 0.1, or 1.0 μM compound in a final concentration of 0.5% (v/v) DMSO. Cell medium was changed every three days by removing half and replacing it with fresh medium containing Aβ42, apoE4, and compound such that the starting concentrations were maintained for the duration of the experiment. At 9 dpe, cells were fixed for immunocytochemistry (ICC) and conditioned medium was collected for analysis of Aβ concentrations by enzyme-linked immunosorbent assay (ELISA). (**B**) Representative ICC images of neurons at 9 dpe labeled for total tau (red), Aβ (white), and cell nuclei (blue). Scale bars = 20 μm. (**C**) Total area of Hoechst^+^ cell nuclei at 9 dpe, relative to the PBS control group. The data represent the mean ± SD of n = 6 wells per group. \**P* < 0.05 by one-way ANOVA. (**D**) Total area of Aβ^+^ pathology at 9 dpe, relative to the PBS control group. The data represent the mean ± SD of n = 6 wells per group. \*\*\**P* < 0.001 by one-way ANOVA. (**E**) Representative ICC images of neurons at 9 dpe to apoE4 and Aβ42 and treated with compounds at 1 μM and labeled for total tau (red), Aβ (white), and cell nuclei (blue). Scale bars = 20 μm. (**F**) Total area of Hoechst^+^ cell nuclei at 9 dpe, relative to the DMSO control group. The data represent the mean ± SD of n = 6 wells for DMSO and n = 3 wells per concentration for compounds. \**P* < 0.05, \*\**P* < 0.01, and \*\*\**P* < 0.001 compared to the DMSO control by one-way ANOVA. (**G**) Total area of Aβ^+^ pathology at 9 dpe, relative to the DMSO control group. The data represent the mean ± SD of n = 6 wells for DMSO and n = 3 wells per concentration for compounds. \**P* < 0.05, \*\**P* < 0.01, and \*\*\**P* < 0.001 compared to the DMSO control by one-way ANOVA.

Each of the eight small molecule hit compounds were dosed into the cell culture medium concurrently with exposure to apoE4+Aβ (Fig. 3A). Six of the compounds had no discernable effects on cell viability or on neuronal morphology at 9 dpe (Fig. 3E). However, two compounds, idarubicin and epirubicin, caused a significant reduction in cell viability at 0.01 µM concentration (Fig. 3F), which is consistent with their clinical use as topoisomerase II inhibitor chemotherapeutics with known toxicity. These compounds may benefit from structural modifications to reduce their side effects while retaining their potent anti-amyloid properties. Sulfacetamide and EGCG produced a slight increase in cell viability at some concentrations, suggesting that they may be neuroprotective. We next examined the effects of the six non-toxic compounds on Aβ neuropathology. At 9 dpe, all compounds exerted a significant effect on Aβ neuropathology at 100 nM and 1 µM, reducing it by 49−71% compared to the DMSO control, and EGCG, olanzapine, and indirubin also reduced Aβ neuropathology at a concentration of 10 nM (Fig. 3G). Additionally, three compounds (i.e., sulfacetamide, EGCG, and olanzapine) significantly reduced the level of Aβ in the conditioned medium at 9 dpe (Fig. S7) which suggests that they may have decreased cellular production or secretion of Aβ.

### Small molecule compounds reduce pTau neuropathology in primary neurons from TgF344 transgenic rats

Aβ induces the phosphorylation and subsequent aggregation of the tau protein into NFT as a key step in the AD pathogenic process (*2*). Because we did not observe tau aggregation in 5xFAD mouse neurons, we turned to the TgF344-AD transgenic rat model that expresses human APP and PSEN1 with familial AD mutations and exhibits robust NFT pathology (*43*). In a similar experimental paradigm as was used for 5xFAD mice (Fig. 4A), primary neurons from TgF344-AD rats exposed to Aβ and apoE4 formed robust intracellular and extracellular Aβ pathology that was accompanied by pTau neuropathology by 14 dpe which included intracellular puncta, axonal blebbing, and neuropil thread-like structures (Fig. S8). Following treatment with each of the five novel and non-toxic hit compounds (i.e., sulfacetamide, imipramine, PD 81723, olanzapine, and indirubin), we observed a significant reduction in the amounts of Aβ neuropathology (Fig. 4B, C), total tau (Fig. 4D), and pTau phosphorylated at the S202/T205 epitope (Fig. 4E) compared to neurons treated with DMSO. Furthermore, PD 81723 and indirubin significantly increased neuronal cell survival (Fig. 4F).

**Fig. 4.**
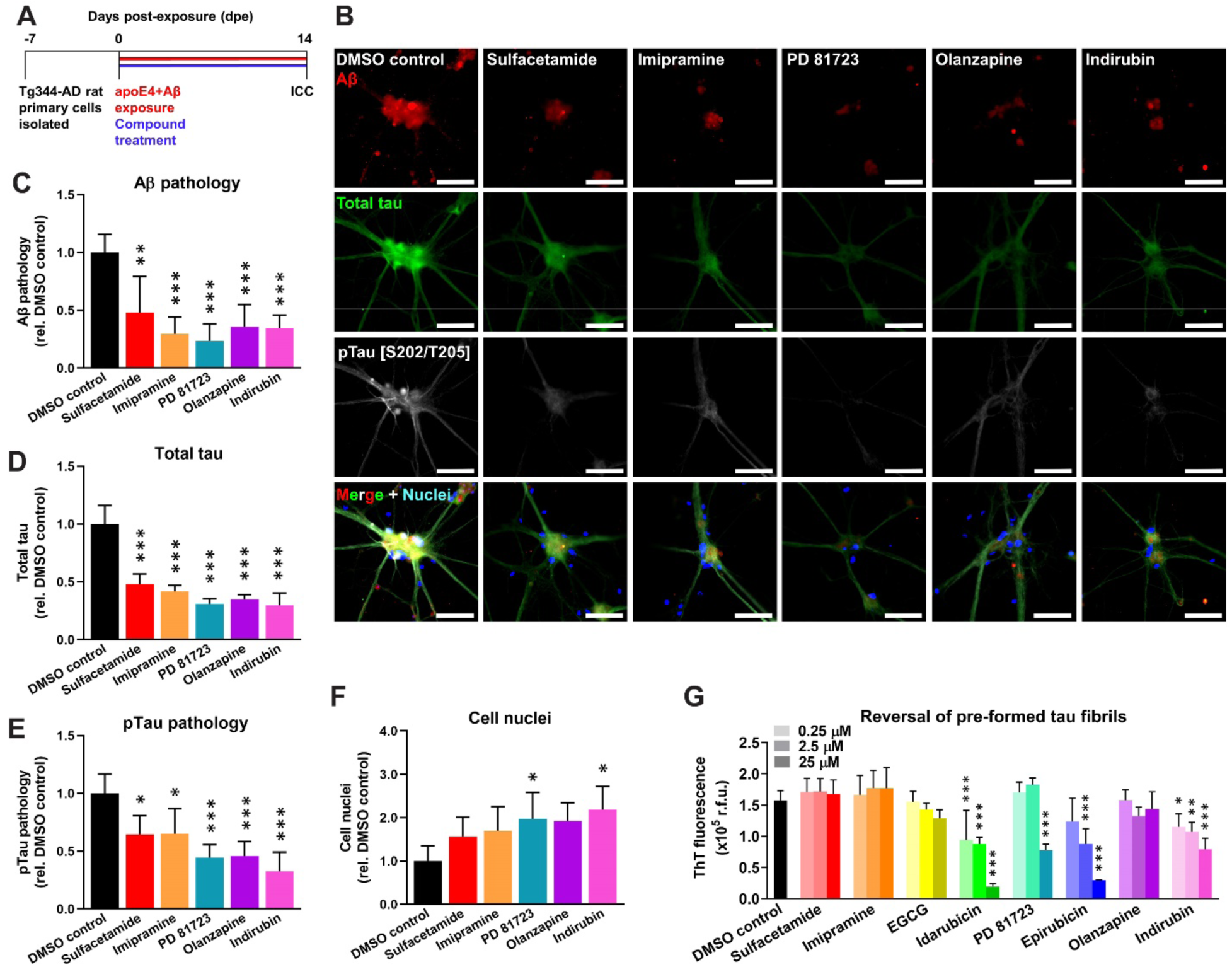
Small molecule compounds inhibit pTau neuropathology in primary neurons from TgF344-AD transgenic rats. (**A**) Schematic of drug efficacy experiments using primary neurons from the TgF344-AD transgenic rat model of AD. One week after cell isolation from day P1 pups, cells were exposed to 100 nM Aβ42 and 1 nM apoE4 and were treated concurrently with 1 μM compound in a final concentration of 0.5% (v/v) DMSO. Cell medium was changed every three days by removing half and replacing it with fresh medium containing Aβ42, apoE4, and compound such that starting concentrations were maintained for the duration of the experiment. At 14 dpe, cells were fixed for ICC. (**B**) Representative ICC images of neurons at 14 dpe, treated with compounds at 1 μM, and labeled for Aβ (red), total tau (green), pTau [S202/T205] (white), and cell nuclei (blue). Scale bars = 50 μm. (**C**) Total area of Aβ^+^ pathology, relative to the DMSO control group. (**D**) Total area of total tau^+^ fluorescence signal, normalized to the total area of Hoechst^+^ fluorescence signal, and relative to the DMSO control group. (**E**) Total area of pTau [S202/T205]^+^ pathology, normalized to the total area of Hoechst^+^ fluorescence signal, and relative to the DMSO control group. (**F**) Total area of Hoechst^+^ cell nuclei at 14 dpe, relative to the DMSO control group. (**C**−**F)** The data represent the mean ± SD of n = 4 wells per group. \**P* < 0.05, \*\**P* < 0.01, and \*\*\**P* < 0.001 compared to the DMSO control by one-way ANOVA. (**G**) Eight hit compounds were tested for disaggregation of pre-formed heparin-induced tau fibrils. The experiment was replicated twice and the results were combined. The data represent the mean ± SD of n = 6 wells per group. \**P* < 0.05, \*\**P* < 0.01, and \*\*\**P* < 0.001 compared to the DMSO control by one-way ANOVA.

To determine whether the hit compounds could act directly on tau, we measured the ability of each compound to disaggregate pre-formed tau fibrils. Idarubicin, PD 81723, epirubicin, and indirubin all significantly reversed tau fibril formation (Fig. 4G). In contrast, sulfacetamide, EGCG, PD 81723, and olanzapine had no effect. Taken together, these data indicate that sulfacetamide, imipramine, and olanzapine reduce pTau neuropathology via inhibition of Aβ, whereas PD 81723 and indirubin act, at least in part, directly on the tau protein.

### Imipramine and olanzapine use correlates with improved outcomes in human AD patients

We next asked whether any of our identified hit compounds were currently being prescribed for other indications, and whether their use was associated with any changes in cognition or risk for developing AD. We acquired longitudinal data from the National Alzheimer’s Coordinating Center (NACC) on 42,661 subjects who were seen at 39 different Alzheimer’s Disease Research Centers (ADRCs) in the U.S. since 2005 (*44*). We searched the prescription drug histories of the subjects in the NACC dataset and found that 40 subjects had taken imipramine, an anti-depressant, and that 94 subjects had taken olanzapine, an anti-psychotic. We then identified ‘control’ subjects who had been prescribed any anti-depressant (n = 6,233 subjects) or any anti-psychotic (n = 798 subjects) medication *other* than imipramine or olanzapine. We first evaluated changes in cognition in all of the subjects over time as measured by the Mini-Mental State Exam (MMSE). Controlling for age and sex, we found that the subjects who took imipramine had a significantly greater change (i.e., improvement) in MMSE score over time compared to subjects who took any other anti-depressant medication (*P* = 0.0490) (Table 2). Likewise, subjects who took olanzapine had a significantly greater change (i.e., improvement) in MMSE score over time compared to subjects who took any other anti-psychotic medication (*P* = 0.0310) (Table 2). Notably, our results show that imipramine use corresponded to an estimated increased score of 0.4186 points (out of 30) per year, and that olanzapine use corresponded to an estimated increased score of 0.4937 points per year, relative to their respective control groups (Table 2). Because we identified imipramine and olanzapine as specific inhibitors of the apoE4-Aβ interaction, we also determined whether *APOE* genotype might influence their effects on cognition. When the subjects who took imipramine were segregated into *APOE4* carriers and *APOE4* non-carriers, both groups showed improvement on imipramine by estimate compared to control, and the estimate for *APOE4* carriers was larger, but none of the contrasts were statistically significant, possibly because the analysis was underpowered due to the small sample size (Table 2). Subjects carrying at least one *APOE4* allele who took olanzapine had a significantly greater change (i.e., improvement) in MMSE score over time (*P* = 0.0235), whereas subjects carrying no *APOE4* allele who took olanzapine showed improved cognition by a lower estimate, and it was not statistically significant (Table 2).

**Table 2.**
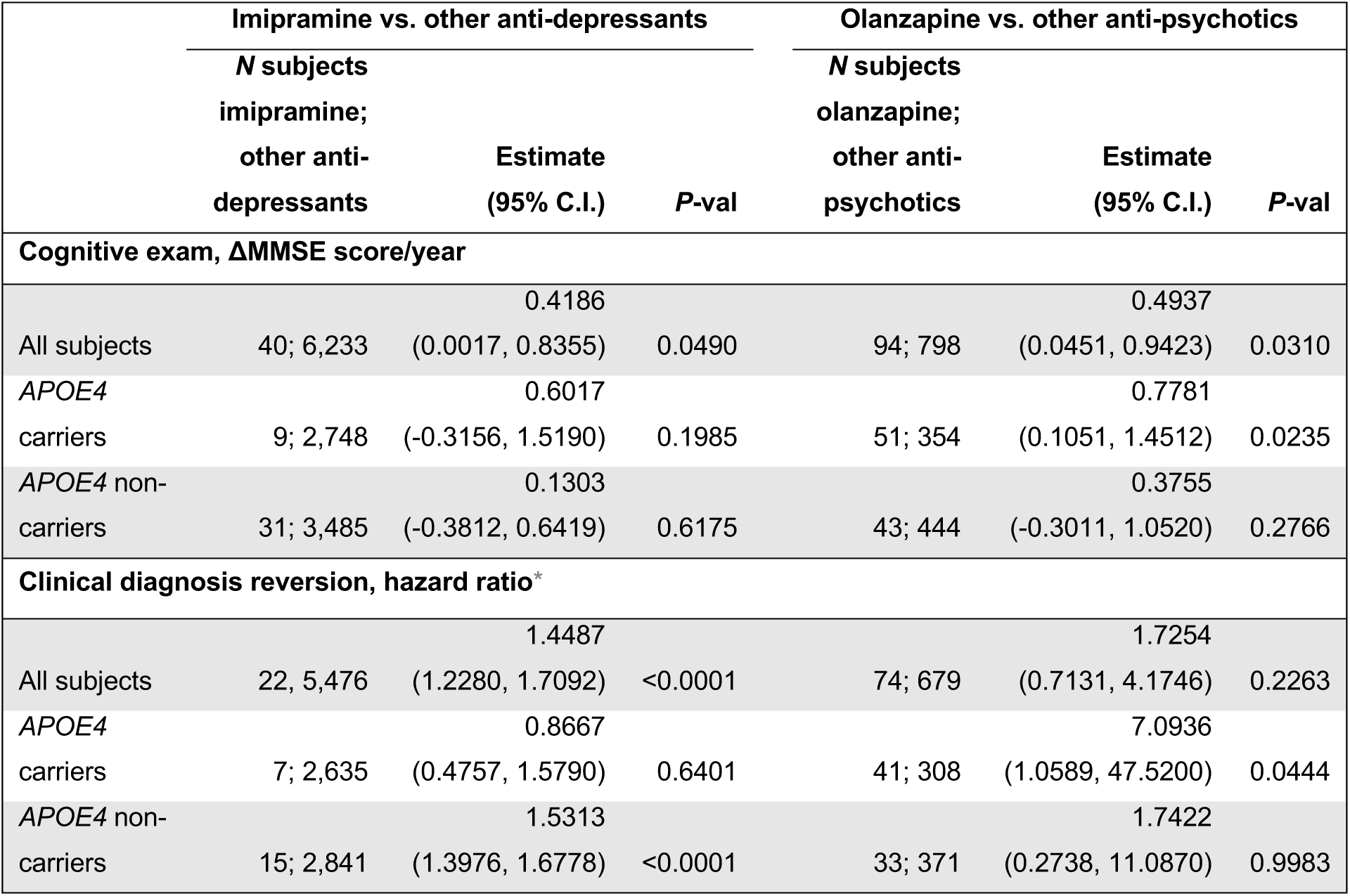
Retrospective analysis of NACC dataset for cognition and clinical diagnosis reversion. The cumulative exposure of imipramine and other anti-depressants were compared and the on/off status of olanzapine and other anti-psychotics were compared, using regression modeling for statistical comparisons of cognitive exam scores and using Cox proportional hazard ratio analysis for statistical comparisons of clinical diagnosis reversion. Test statistics and degrees of freedom are provided in Data file S3. *Only subjects who reported use of a medication prior to clinical diagnosis reversion were included.

|  | Imipramine vs. other anti-depressants |  |  | Olanzapine vs. other anti-psychotics |  |  |
| --- | --- | --- | --- | --- | --- | --- |
|  | *N* subjects<br>imipramine;<br>other anti-depressants | Estimate<br>(95% C.I.) | *P*-val | *N* subjects<br>olanzapine;<br>other anti-psychotics | Estimate<br>(95% C.I.) | *P*-val |
| **Cognitive exam, ΔMMSE score/year** |  |  |  |  |  |  |
|  |  | 0.4186 |  |  | 0.4937 |  |
| All subjects | 40; 6,233 | (0.0017, 0.8355) | 0.0490 | 94; 798 | (0.0451, 0.9423) | 0.0310 |
| *APOE4* |  | 0.6017 |  |  | 0.7781 |  |
| carriers | 9; 2,748 | (-0.3156, 1.5190) | 0.1985 | 51; 354 | (0.1051, 1.4512) | 0.0235 |
| *APOE4* non- |  | 0.1303 |  |  | 0.3755 |  |
| carriers | 31; 3,485 | (-0.3812, 0.6419) | 0.6175 | 43; 444 | (-0.3011, 1.0520) | 0.2766 |
| **Clinical diagnosis reversion, hazard ratio\*** |  |  |  |  |  |  |
|  |  | 1.4487 |  |  | 1.7254 |  |
| All subjects | 22, 5,476 | (1.2280, 1.7092) | <0.0001 | 74; 679 | (0.7131, 4.1746) | 0.2263 |
| *APOE4* |  | 0.8667 |  |  | 7.0936 |  |
| carriers | 7; 2,635 | (0.4757, 1.5790) | 0.6401 | 41; 308 | (1.0589, 47.5200) | 0.0444 |
| *APOE4* non- |  | 1.5313 |  |  | 1.7422 |  |
| carriers | 15; 2,841 | (1.3976, 1.6778) | <0.0001 | 33; 371 | (0.2738, 11.0870) | 0.9983 |

We next determined whether subjects received an improved clinical diagnosis from their physician after taking imipramine or olanzapine. We used Cox Proportional Hazard models to evaluate the incidence of a subject reverting from a clinical diagnosis of AD to mild cognitive impairment (MCI), or of reverting from a diagnosis of MCI to normal cognition (NC). Controlling for age and sex, we found that, compared to subjects who took any other anti-depressant medication, subjects who took imipramine had increased incidence of reverting toward normal in clinical diagnosis by an estimated 44.87% for each additional year of exposure, a result that was highly statistically significant (*P* < 0.0001) (Table 2). *APOE4* carriers who took imipramine also had significantly decreased incidence of worsening their clinical diagnosis (from NC to MCI or from MCI to AD) compared to *APOE4* non-carriers (*P* = 0.0474) (Table S1). Given that other anti-depressants have been previously proposed as AD therapeutics, particularly selective serotonin reuptake inhibitors (SSRIs) (*45, 46*), we also directly compared imipramine to several other common anti-depressants. The incidence of reverting toward normal in clinical diagnosis was significantly higher for imipramine compared to two common SSRIs, fluoxetine and citalopram, and compared to doxepin, a tricyclic anti-depressant with similar pharmacological properties to those of imipramine (Fig. S9A).

The incidence of reverting toward normal in clinical diagnosis when taking olanzapine was an estimated 72.54% greater compared to control anti-psychotic medications, although this result was not statistically significant (Table 2). Among *APOE4* carriers, being on olanzapine increased the incidence of reverting in clinical diagnosis by an estimated factor of 4.8064 compared to control (*P* = 0.0444). The incidence of reverting toward normal while taking olanzapine was not significantly different from the common anti-psychotics aripiprazole and quetiapine (Fig. S9B). Interestingly, aripiprazole, which showed the greatest trend toward increased incidence of clinical diagnosis reversion, was a hit in our exploratory drug screen (Data file S2), although it did not produce a dose-dependent response in our HTS assay (Fig. 2A) and we have not pursued it further.

Finally, we evaluated the relationship between sex and age and the potential effects of imipramine or olanzapine on clinical diagnosis compared to controls. We found that cumulative imipramine exposure significantly increased the incidence of reverting in clinical diagnosis for men between the ages of 66.5 and 88.5 years, although the effect was not statistically significant in women (Fig. S9C). Olanzapine showed a trend toward greater benefit for subjects of older age, although the result was not statistically significant (Fig. S9D). Taken together, these results indicate that, compared to other anti-depressant and anti-psychotic medications, the ability of imipramine and olanzapine to specifically inhibit apoE-catalyzed Aβ fibrillization predicts their specific ability to improve cognition and reverse clinical diagnosis toward normal.

## Discussion

Since the development of ThT-based amyloid fibrillization assays in the 1980s, a wide range of concentrations and assay conditions have been evaluated with no clear consensus (*47*).

Therefore, our first objective was to determine the optimal conditions of a ThT-based assay for studying the effects of apoE on Aβ fibril formation. We employed DOE, a statistical method for process optimization that allows experimentation on numerous variables at the same time, each at a wide range of values. In contrast to traditional “one variable at a time” methods, DOE is highly efficient and also identifies relevant interactions between variables. As an example of its efficiency, in our first response surface experiment, we evaluated five different Aβ, apoE4, and ThT concentrations each using 32 combinations, rather than testing all 5 x 5 x 5 = 125 possible combinations. In several experiments, each building on the previous one, we ultimately identified 20.9 µM Aβ42, 0.75 nM apoE4, and 14.8 µM ThT as the optimal concentrations that maximize Aβ fibril formation in PBS at 37°C.

We identified an optimal concentration of 0.75 nM apoE4, which was surprising, given that a much higher concentration of apoE4 was used in the first studies demonstrating that apoE4 accelerates Aβ fibril formation, although other concentrations were not tested (*15, 16*). The physiological concentration of apoE in human plasma is 4 µM, and is 100 nM in human cerebrospinal fluid (CSF) (*48, 49*). However, it is important to consider that apoE and Aβ are most likely to interact in the brain interstitial fluid (ISF), especially around synapses where Aβ is produced and exerts its neurotoxic effects (*50*). Thus, the most relevant benchmark may be the apoE concentration in brain ISF, which has been shown to be 0.30 nM for wild-type mice and 0.37 nM for human *APOE4* knock-in (KI), measured by *in vivo* microdialysis (*51*). When 5xFAD mice were crossed with the same *APOE4* KI mice, they showed significantly accelerated plaque deposition (*52*), suggesting that apoE concentrations similar to those that were tested here are sufficient to catalyze Aβ fibrillization *in situ*. With respect to the low apoE4:Aβ ratio used in our experiments, it should be noted that apoE has two binding sites for Aβ (*53*), and that Aβ42 often exists in a polymeric β-sheet structure in the AD brain, for which apoE has greater affinity (*54*). Based on the traditional definition of a catalyst, we have also hypothesized that apoE may not be consumed in the catalytic reaction, but may instead be released from Aβ and thereby catalyze the formation of multiple fibrils (*55*), although this has yet to be demonstrated. In this case, very low concentrations of apoE4 may exert a significant amyloidogenic effect, underscoring the importance of inhibiting its interaction with Aβ. Our results also indicate that the catalytic effect of apoE is highly dependent upon the apoE:Aβ molar ratio, which may explain, in part, conflicting reports on its amyloidogenic effects *in vitro* (*56*).

Our HTS assay identified eight compounds with potent activity against Aβ aggregation or against the catalytic effect of apoE4 on Aβ fibrillization. Interestingly, our reversal studies then showed that five of those compounds — EGCG, idarubicin, PD 81723, epirubicin, and indirubin — disaggregated pre-formed fibrils of Aβ (Fig. 2B−E), which suggests that these molecules may be pursued as interventional treatments for patients with pre-existing AD neuropathology. On the other hand, we found that sulfacetamide, imipramine, and olanzapine did not block or reverse Aβ fibrillization independent of apoE4, suggesting that they are specific inhibitors of the apoE4-Aβ interaction and warrant further development for preventing AD, particularly in *APOE4* carriers. We then tested all eight hit compounds in primary neurons from 5xFAD mice that overproduce human Aβ leading to both intraneuronal and extracellular Aβ neuropathology (*42*), which was accelerated by the addition of human apoE4 and Aβ42 to the culture medium as seeds. ApoE is capable of penetrating the cell membrane and enhancing neuronal Aβ uptake (*57, 58*), and may have thereby contributed to the intraneuronal aggregation of Aβ. Human apoE4 itself has been reported to be toxic to neurons in culture and in mice (*57*), but we did not observe this effect at the very low concentration used. Therefore, we believe that the effects of apoE4 in our neuronal assays were predominantly via catalysis of Aβ fibril formation. Despite using isolation and culture methods that favored neurons, sparse astrocytes, which secrete apoE, are often present in neuronal cultures, and neurons also secrete apoE under stressed conditions (*59*). Thus, we cannot rule out a potential contribution of mouse- or rat-derived apoE to the Aβ and pTau neuropathologies observed in our cellular assays. We found that all six non-toxic hit compounds reduced Aβ pathology in 5xFAD mouse neurons (Fig. 3G), either by inhibiting the effect of apoE4, or by preventing/reversing Aβ aggregation, and we then confirmed this effect in a second model using Tg344 rat primary neurons (Fig. 4C). Importantly, we also showed that the compounds reduced the subsequent intraneuronal accumulation of pTau protein (Fig. 5D and E), which is directly linked to neurodegeneration and cognitive decline (*3, 60*). The effects of PD 81723 and indirubin on pTau pathology may have been, in part, via direct action on tau oligomers/fibrils (Fig. 4G). However, sulfacetamide, imipramine, and olanzapine showed no direct effects on tau fibrils, indicating that they may reduce pTau pathology and subsequent neurodegeneration indirectly via inhibition of the effect of apoE4 on Aβ.

Imipramine is a tricyclic antidepressant that blocks norepinephrine and serotonin reuptake. Given the frequent use of antidepressants by AD patients, imipramine has also been evaluated in cell and mouse models of AD where it was found to reduce Aβ accumulation (*61, 62*). Olanzapine is an anti-psychotic drug that has been evaluated for acute treatment of behavioral and psychological symptoms of AD (*63*). Olanzapine has not been tested clinically as a disease-modifying therapy for AD, but it has been shown to have neuroprotective effects against Aβ-induced oxidative stress and apoptosis (*64, 65*). Indirubin, a natural compound, was found to both prevent and to reverse Aβ fibrillization in our studies. Indirubin is best known for being a potent inhibitor of cyclin-dependent kinases (CDKs) and glycogen synthase kinase-3β (GSK-3β), both of which phosphorylate tau. Therefore, indirubin may have multi-functional therapeutic benefits for AD. Indeed, indirubin has been reported to reduce amyloid and tau pathology, attenuate neuroinflammation, and improve spatial memory deficits in AD mouse models (*66*). Imipramine, olanzapine, and indirubin demonstrated efficacy in our cellular Aβ assay at nanomolar concentrations (Fig. 3H). These compounds are exceptionally promising because they may accommodate peripheral dosing, for which CNS bioavailability is very low, even for BBB-permeable drugs. Maintaining a therapeutic drug concentration in the brain is crucial because inhibiting peripheral apoE or increasing its levels by parabiosis have been shown to have no effect on Aβ deposition in the brain (*67, 68*). We also identified sulfacetamide, an antibiotic, and PD 81723, an allosteric enhancer of brain adenosine A1 receptors, as novel therapeutic candidates that have not been previously evaluated for AD.

Depression and psychosis are well-known co-morbidities of AD and other dementias. As such, a significant proportion of NACC participants reported use of anti-depressant and/or anti-psychotic medications, providing large control populations with similar clinical presentations that enabled us to evaluate the potential effects of imipramine and olanzapine. Our analyses show that, compared to these control populations, subjects taking imipramine or olanzapine had improved cognition and diagnoses, direct clinical measures of disease severity. Notably, in our drug screen, we found that imipramine and olanzapine strongly inhibited the apoE4-catalyzed fibrillization of Aβ, whereas none of the other anti-depressants or anti-psychotics in the NCC library had any such activity. In line with our identified mechanism of action, these apoE4 inhibitors also demonstrated a preferential benefit for *APOE4* carriers over non-carriers, with those taking olanzapine having a greater change (i.e., improvement) in MMSE score (Table 2), and those taking imipramine having reduced incidence of clinical diagnosis conversion (Table S1). Furthermore, cumulative imipramine exposure was associated with significantly greater incidence of clinical diagnosis reversion towards normal compared to fluoxetine and citalopram (Fig. S9), two SSRIs proposed to reduce Aβ production via increased serotonin signaling that have been evaluated in humans (*69*). Taken together, these results provide strong evidence of the potential clinical benefits of imipramine and olanzapine use in human subjects and support further development and evaluation of these and our other hit compounds as disease-modifying treatments for AD.

The clinical diagnoses recorded in the NACC database were frequently made in consensus conferences, wherein at least one physician and one neuropsychologist evaluated a subject’s MMSE score, neuropsychological exam, and full clinical history, among other information (*44*). The fact that we found both imipramine and olanzapine use to be associated with improvements in clinical diagnosis, for which MMSE (a memory-focused exam) was weighed only in part, suggests that these drugs may have had additional functional benefits not identified here, but which were taken into account in the consensus conferences. It is also possible that beneficial neuropsychological effects of imipramine and olanzapine, via their primary mechanisms of action, contributed to the improvements observed in our study. However, imipramine and olanzapine have not been found to be particularly effective for treating depression or psychosis in AD patients (*63, 70*). Both drugs have known interactions and side effects and are prescribed cautiously in elderly patients (*71*), which is likely a reflection of the dosages necessary to achieve their anti-depressant or anti-psychotic effects. Future prospective clinical studies are warranted to identify the appropriate dosing for different age groups that avoids these known side effects while maintaining the therapeutic potential for prevention/reversal of AD.

Novel CNS drug development has historically been a 10- to 17-year-long process with less than a 10% chance of success and a cost of approximately $1.8 billion per drug (*72*). For drugs targeting AD, novel drug development has been especially challenging due to the slow progression of disease requiring lengthy clinical trials with a large number of participants, and due to the lack of robust and predictive biomarkers (*4*). Although a number of drugs targeting Aβ are currently being tested, there has been a very high failure rate of ∼99.6% for AD therapeutics in clinical trials, and there are currently no approved disease-modifying treatments for AD (*4*). Drug repurposing, using known drugs for novel indications, has several unique advantages for AD. There is existing knowledge from prior clinical trials on the pharmacological effects, pharmacokinetics, toxicology, and side effects in humans. Therefore, drugs with good safety profiles can be prioritized, expediting the early phases of clinical testing and reducing the failure rate. Development costs are significantly less for repurposed drugs, increasing the chances that a company will be willing to invest to bring a drug to market. For these reasons, a repurposed drug in Phase II trials has greater than twice the likelihood of making it to market than a novel drug (*73*). Indeed, one of the most widely prescribed medications to reduce some symptoms of dementia is memantine, which was originally developed as an anti-viral drug and was then serendipitously found to have anti-glutamatergic activity and was repurposed for AD (*74*). Other high-content phenotypic screens have been developed in recent years aimed at drug repurposing for AD (*75*); however, none have focused on inhibition of apoE as a key driver of disease. For the reasons highlighted above, the methods used here to identify drug candidates with some safety/dosing information available, good BBB permeability, and strong preclinical efficacy positions them well to reach the clinic as disease-modifying therapeutics for AD.

These biochemical, cellular, and clinical results strongly support the concept that apoE serves as a catalyst for fibrillization of Aβ into neurotoxic oligo/polymers and that further studies on this approach to the development of AD therapeutics are warranted. Furthermore, apoE has been implicated in a number of Aβ-independent pathogenic mechanisms that cause Parkinson’s disease, primary tauopathies, and amyotrophic lateral sclerosis, among other disorders (*76*). Thus, the apoE-centric screening methods and drug candidates we report here may also prove valuable for addressing other human neurodegenerative diseases.

## Materials and Methods

### Study design

The objective of this study was to develop a HTS assay, employ it to identify novel AD therapeutics from a drug library, and confirm their efficacy in a primary neuron model of AD. In assay optimization experiments, 3−8 wells were used per group and experiments were replicated one or two times, as indicated in the figure legends. When replicated twice, experiments were performed on different days and in different plates and the results of the two experiments were combined. For HTS, each compound was tested in 3−4 wells per concentration and experiments were replicated one or two times, as indicated in the figure legends. For the primary neuron models, we used the minimum number of mice and rats to obtain sufficient numbers of cells to test all compounds in three wells per concentration. Cells from individual mice were pooled and used for all groups to remove the effect of biological variation and to allow us to use fewer mice. Cells from individual rat pups were not pooled in order to evaluate the drug effects on different biological replicates, although each drug and controls were tested on cells derived from the same rats.

### Development of the apoE4-Aβ fibrillization assay

Recombinant human Aβ42 sodium hydroxide (NaOH) salt (rPeptide) was received following pre-treatment to ensure a consistent monomeric preparation, as described previously (*77*). For NaOH pre-treatment of Aβ peptide, briefly, following recombinant protein expression and purification, Aβ42 peptides were dissolved in 2 mM NaOH, pH 10.5, and then sonicated and lyophilized. Upon receipt, the lyophilized peptide was reconstituted in ice-cold Dulbecco’s phosphate buffered saline (DPBS), pH 7.4, which avoids the solution passing through the isoelectric point of Aβ (pI = 5.5), which would induce aggregation (*78*). The reconstituted Aβ42 stock solution was quickly aliquoted and snap-frozen in liquid nitrogen and then stored at −80°C until use. Great care was taken to ensure consistency and reproducibility across all experiments by using Aβ from a single batch, thawing and maintaining Aβ stock on ice until use, and never re-freezing the unused portion of thawed Aβ stock. For fibrillization experiments, Aβ42, recombinant human apoE4 (Sigma), and ThT (Sigma) were combined at pre-determined concentrations in DPBS in a total volume of 40 µl in a 384-well µ-clear bottomed plate (Greiner). Plates were sealed to prevent evaporation and incubated at 37°C with constant rapid agitation and the fluorescence intensity of ThT at λ_ex_= 440 nm, λ_em_= 490 nm was measured every 10 min for up to 24 h using a fluorescence plate reader (Biotek). Once the optimal concentrations of approximately 20 μM Aβ42, 1 nM apoE, and 15 μM ThT were determined, they were maintained throughout subsequent studies unless noted otherwise. For HTS assay validation, recombinant human Aβ40 (rPeptide), recombinant human scrambled Aβ42 (rPeptide), recombinant human apoE2 and apoE3 (Creative Biomart), recombinant human apoA-I (Creative Biomart), human plasma-derived apoE (Sigma), or DMSO (Sigma) were included or substituted at the indicated concentrations.

### HTS of the NCC library

The NCC library was developed by the National Center for Advancing Translational Sciences (NCATS) (https://ncats.nih.gov/smr). Detailed information about these compounds is available using the NIH Chemical Genomics Center (NCGC) Pharmaceutical Collection browser (*79*). The NCC library was received from Evotec, Inc. and contained each compound at 10 mM in DMSO which were aliquoted and stored at −80°C until use. To set up the exploratory drug screen, compounds were thawed, diluted in DMSO, and added at a concentration of 2 µM to Aβ42 (2 μM) in water, followed by the addition of apoE4 (20 nM) and the mixture was incubated at rt for 15 min. The mixture was then divided into 3 separate wells of a 96-well plate, ThT (8 μM) and glycine (30 mM) were added for a total volume of 125 μL per well and the plate was incubated at rt for 10 min in the dark. The fluorescence intensity of ThT was then measured using the fluorescence plate reader. The 595 compounds were divided across numerous plates, and compounds on each plate were compared to control wells on the same plate that received Aβ42, apoE4, ThT, and DMSO. Unlike in the exploratory screen, the optimal concentrations of Aβ42, apoE4, and ThT described above were used in 384-well plates in the HTS assay. To set up the HTS assay, compounds were thawed, diluted in DMSO, and added to the Aβ42 in DPBS at final concentrations of 0.25, 2.5, and 25 µM in 5% DMSO/ 95% DPBS (v/v), followed immediately by the addition of apoE4 and ThT in a total volume of 40 μL per well. Plates were sealed to prevent evaporation and incubated at 37°C with constant shaking and the fluorescence intensity of ThT was measured every 10 min for 24 h using the fluorescence plate reader. The 87 compounds were divided across three separate plates, and compounds on each plate were compared to control wells on the same plate that received Aβ42, apoE4, ThT, and 5% DMSO. The criteria for hit identification were that the compound reduced ThT fluorescence by at least 30% at any concentration and that the effect was generally dose-dependent.

### Inhibition of Aβ alone and disaggregation of pre-formed fibrils

Each of the eight hit compounds was added to Aβ42 at 0.25, 2.5, and 25 µM in 5% DMSO/ 95% DPBS (v/v), followed immediately by the addition of ThT and measurement of fluorescence intensity every 10 min for 24 h. The AUC of ThT fluorescence intensity was calculated and normalized to control wells receiving Aβ42, ThT, and 5% DMSO. To test compounds for disaggregation of pre-formed Aβ fibrils, Aβ42 and apoE4 were combined and incubated at 37°C for 24 h with constant shaking to induce fibrillization. Pre-formed Aβ fibrils were then divided into separate wells, and compounds were added in a final concentration of 5% DMSO and incubated at rt for 30 min with constant shaking. ThT was added to each well, the plates were incubated at rt for 15 min, and then fluorescence intensity was measured and normalized to control wells receiving only 5% DMSO. To test compounds for disaggregation of pre-formed tau fibrils, 2 μM recombinant human K18 tau peptide (Novus), comprising the microtubule binding domain of the 4R tau isoform, was combined with 2 μM heparin (Sigma) and 300 μM dithiothreitol (DTT, Invitrogen) in DPBS and incubated at 37°C for 24 h with constant shaking to induce fibrillization. Pre-formed tau fibrils were then divided into separate wells, and compounds were added in a final concentration of 5% DMSO and incubated at rt for 30 min with constant shaking. ThT (12.5 μM) was added to each well, the plates were incubated at rt for 15 min, and then fluorescence intensity was measured and normalized to control wells receiving only 5% DMSO.

### Transmission electron microscopy

Immediately following the measurement of ThT fluorescence intensity, pre-formed Aβ fibrils treated with individual hit compounds, or with DMSO as a control, were applied undiluted to Formvar/carbon-coated copper grids with 300 square mesh (Electron Microscopy Sciences) for 2 min. Grids were gently blotted on filter paper (Whatman) to remove excess fibrils, then washed twice in water and stained with 2% (w/v) uranyl acetate (Electron Microscopy Services) twice for 20 s each, blotting on filter paper in between each step. Grids were air dried and imaged on a Tecnai G^2^ Spirit BioTwin microscope (FEI) at 80 kV with a side-mount digital camera (AMT Imaging). TEM images were processed and analyzed using Fiji version 2.1.0/1.53c.

### Animals

5xFAD transgenic mice, which express human APP harboring the Swedish (K670N/M671L), Florida (I716V), and London (V717I) familial AD mutations, and human PSEN1 harboring the M146L and L286V familial AD mutations, from two separate transgenes, each driven by the murine Thy1 promoter, were originally developed on a mixed B6SJL background (*42*). 5xFAD mice that had been backcrossed to a congenic C57BL/6J background (Jackson Labs # 034848-JAX) were received and maintained as a hemizygous line by breeding with C57BL/6J mice. TgF344-AD transgenic rats, which express human APP harboring the Swedish (K670N/M671L) and human PSEN1 with the Δ exon 9 mutation, both driven by the mouse prion protein (PrP) promoter (*43*), were maintained on a Fischer 344 background Mice and rats were treated in accordance with the *Guide for the Care and Use of Laboratory Animals*. All procedures were approved by the Institutional Animal Care and Use Committee of the University of Colorado.

### 5xFAD mouse primary neuron cell model

P1-P2 5xFAD mouse pups were genotyped using primer probes and real-time polymerase chain reaction (RT-PCR) analysis of 1 mm tail snip samples. Brains from the mouse pups were then rapidly removed, cerebral cortices were isolated using a sterile razor blade, and tissue from multiple pups were pooled for experiments. Primary cultures of neurons were prepared using the Papain Dissociation System (Worthington) according to the manufacturer’s instructions. To prepare neuronal cultures, cortical tissue was dissociated in 20 U/ml papain under constant agitation at 37°C for 45 min. A single cell suspension was obtained by trituration, then papain was inactivated using ovomucoid protease inhibitor and cells were filtered through a 100 μm cell strainer and diluted in warm Neurobasal medium supplemented with Glutamax, B27 supplement, and penicillin/streptomycin (all from Gibco). Cells were seeded at 30,000 cells/cm^2^ in 96-well μ-clear bottomed plates (Ibidi) pre-coated with 10 μg/ml poly-D-lysine (Sigma). Neural cultures were maintained at 37°C in a humidified 5% CO2 chamber for 3 d, and then half of the culture medium was replaced with fresh medium also containing CultureOne supplement (Thermo Fisher), which reduces glial cell proliferation to favor neuronal culture. Every 3 d following exposure to Aβ and apoE4 and treatment with compounds, half of the culture medium was replaced and Aβ42, apoE4, and the test compounds in DMSO were added to maintain the initial concentrations. After 7 d in culture, half of the culture medium was replaced, and 100 nM Aβ42 and 1 nM apoE4 were added, followed by the addition of test compounds at 0.01, 0.1, or 1 μM in a final concentration of 0.5% (v/v) DMSO. At 9 dpe wells were fixed for immunocytochemistry and the conditioned media was collected for ELISA analysis. We used the minimum number of mice to obtain sufficient numbers of cells to test all compounds in three wells per concentration. Cells from individual mice were pooled and used for all groups to remove the effect of biological variation and to allow us to use fewer mice.

### Immunocytochemistry

At the pre-determined end points, the culture medium was removed, and the cells were washed once with DPBS, fixed in 4% (w/v) paraformaldehyde for 30 min, washed four times with DPBS, and stored at 4°C. The cells were permeabilized with 0.1% Triton X-100 in DPBS for 10 min and then blocked with 3% bovine serum albumin (BSA) in DPBS for 90 min and then incubated overnight at 4°C with primary antibodies in 3% BSA in DPBS. 5xFAD mouse cells were labeled with chicken anti-tau (PhosphoSolutions, 1:1000) and mouse anti-Aβ (82E1, IBL, 1:500). TgF344 rat cells were labeled with chicken anti-tau (PhosphoSolutions, 1:1,000), rabbit anti-Aβ (OC, Millipore, 1:500), and mouse anti-pTau (AT8, Sigma, 1:250). Cells were washed and then incubated with Alexa Fluor Plus-conjugated secondary antibodies (Thermo Fisher, 1:500) for 45 min at rt in 3% BSA in DPBS. Cells were washed, and then nuclei were stained with 1 µg/ml Hoechst 33342 (Thermo Fisher) in DPBS for 10 min. The cells were then washed and imaged on an Olympus IX83 inverted fluorescence microscope. Images of entire wells were captured at 20X magnification and then analyzed using Cell Sens v1.12 software (Olympus).

### Enzyme-linked immunosorbent assay (ELISA)

Aβ concentration in conditioned medium from individual wells measured using the human Aβ42 ELISA kit (Thermo Fisher), following the manufacturer’s instructions. Two technical replicates were performed in the ELISA assay for each of three different wells per compound per concentration.

### TgF344-AD rat primary neuron cell model

Brains from P1 TgF344-AD transgenic rat pups were removed, and cortices were isolated using a sterile razor blade. Primary cultures of neurons were prepared from cerebral cortices using the Papain Dissociation System (Worthington) according to the manufacturer’s instructions and were plated and cultured as described above for 5xFAD mouse neurons. Cells from individual rat pups were not pooled but were cultured in separate wells. After 7 d in culture, half of the culture medium was replaced, and 100 nM Aβ42 and 1 nM apoE4 were added, followed by the addition of test compounds at 1 μM in a final concentration of 0.5% (v/v) DMSO. Every 3 d thereafter, half of the culture medium was replaced and Aβ42, apoE4, and the test compounds in DMSO were added to maintain the initial concentrations. At 14 dpe the cells were fixed for immunocytochemistry. We used the minimum number of rats to obtain sufficient numbers of cells to test all compounds in three wells per concentration. Cells from individual rat pups were not pooled in order to evaluate the drug effects on different biological replicates, although each drug and controls were tested on cells derived from the same rats.

### NACC data analysis

The NACC UDS v3 (*44*) was received on April 17, 2020 and contained standardized longitudinal clinical data on 42,661 subjects seen at ADRCs beginning in September, 2005 thru the March, 2020 data freeze. Subjects who had reported taking at least one of the eight hit compounds (sulfacetamide, imipramine, EGCG, idarubicin, PD 81723, epirubicin, olanzapine, or indirubin) were identified by searching the ‘DRUGS’ column. Only subjects with at least two clinic visits and who had reported taking a medication prior to their final clinic visit were considered. Control groups of subjects taking anti-depressant medications or anti-psychotic medications were identified using the ‘NACCADEP’ or ‘NACCAPSY’ columns, respectively, with subjects who reported only taking imipramine or olanzapine being removed. The groups partially overlapped, as, for example, a subject may have reported using imipramine or olanzapine and then reported using a different anti-depressant or anti-psychotic. In developing the models, medication was treated as a time-varying explanatory variable in order to accurately model exposure, as subjects’ medication statuses changed over time. When a medication was listed at a given time point, the exposure was assumed to have been started at the mid-point between the current and previous time points, and to have lasted until the mid-point between the current and subsequent time points. The mean change in MMSE score (ΔMMSE) over time was modeled using time slopes, with time-varying drug and covariate interactions as slope modifiers. Longitudinal regression models were developed using a random time slope by subject and a continuous 1^st^ order auto-regressive covariance structure for errors on the same subjects, and were fit using MMSE scores extracted from the ‘NACCMMSE’ column, and using subjects’ age and sex, identified in the ‘NACCAGE’ and ‘SEX’ columns, as covariates. Due to limited sample sizes, only two-way interactions were considered, and linear effects were assumed. Central limit theorems protect against non-severe departures from nomality, and MMSE is a validated scale. *APOE* models were also developed using the presence or absence of an *APOE4* allele as a modifier of the drug effect. The ‘NACCNE4S’ column was evaluated and subjects with a 1 or 2 were designated *APOE4* carriers, subjects with a 0 were designated *APOE4* non-carriers, and subjects with a 9 (missing data) were excluded. Linear combinations of parameters were tested with T and F tests, and the Satterthwaite method was used to calculate the denominator degrees of freedom. All tests were two-sided, and 95% confidence intervals were presented for all univariate contrasts. For reversion and conversion models, Cox proportional hazards models were developed, stratified by clinical diagnosis. The Cox model makes no parametric assumptions about the shape of the underlying hazard function, and stratification permits different underlying hazard functions for different clinical diagnoses. Tests for violation of the proportional hazards assumptions are not available for models with time-varying covariates. Clinical diagnoses were extracted from the ‘NACCUDSD’ column, in which a 1 was considered “NC”, 2 (cognitively impaired, but not meeting the classical definition for MCI) or 3 were considered “MCI”, and 4 was considered “AD”. A 4 in the ‘NACCUDSD’ column indicates a diagnosis of dementia, which may include AD, Lewy body dementia, frontotemporal dementia, etc. However, the ‘NACCALZD’ column indicated that the vast majority of subjects receiving a dementia diagnosis were deemed to be of AD etiology (e.g., 29/32 subjects who took imipramine), and thus herein, we refer to this group collectively as AD patients. Drug exposure was modeled using time-varying covariates and cumulative exposure, controlled for time since last exposure, was selected for anti-depressants, while on/off status was selected for anti-psychotics. The variance calculations accounted for repeated measures, as the subjects could have multiple reversion/conversion events. Subjects with an initial clinical diagnosis of AD were excluded from the risk set for conversion, and subjects with an initial clinical diagnosis of NC were excluded from the risk set for reversion, as they were ineligible for the event. Only subjects who reported use of a medication prior to a clinical diagnosis reversion or conversion were included in those groups. Age and sex were controlled for, and in the interaction models, all two-way interactions between drug exposure, age, and sex were considered. For the MMSE models, effects were assumed to be linear and only two-way interactison were considered due to limited sample sizes. Linear combinations of parameters were tested with Z and Χ^2^ tests. Hazard was modeled on a logarithimic scale and then the results were transformed back to hazard ratios. All tests were two-sided and 95% confidence intervals were presented for univariate contrasts. For the mixed medications models, subjects taking doxepin, citalopram, fluoxetine, aripiprazole, or quetiapine were identified by searching the ‘DRUGS’ column, and reversion models were developed as described above. Summary statistics including baseline age, sex, baseline MMSE score, drug exposure time, number of clinical diagnosis reversions, number of clinical diagnosis conversions, and test statistics and degrees of freedom for all analyses are provided in Data file S3. Multiple testing adjustment was not applied because of the limited power and exploratory nature of the study.

### Statistical analysis

DOE and statistical analyses for the development of the fibrillization assay were performed using Minitab 18. Linear regression and one-way analysis of variance (ANOVA) were performed using GraphPad Prism 8. Following ANOVA, comparisons between multiple groups was done by post-hoc testing using the Holm-Šidák method and a *P* < 0.05 was considered statistically significant.

## Acknowledgments

We thank L. Johnson, MS (JMC Data Experts, Pittsburgh, PA) for assistance with the DOE and statistical analysis, A. Goldblach (University of Chicago, Chicago, IL) for assistance with the exploratory drug screen, and P. Pressman, MD (University of Colorado Anschutz Medical Campus, Aurora, CO) for helpful discussions on the clinical implications of our NACC data analysis findings. The NACC database is funded by NIA/NIH Grant U01 AG016976. NACC data are contributed by the NIA-funded ADRCs: P30 AG019610 (PI Eric Reiman, MD), P30 AG013846 (PI Neil Kowall, MD), P30 AG062428-01 (PI James Leverenz, MD) P50 AG008702 (PI Scott Small, MD), P50 AG025688 (PI Allan Levey, MD, PhD), P50 AG047266 (PI Todd Golde, MD, PhD), P30 AG010133 (PI Andrew Saykin, PsyD), P50 AG005146 (PI Marilyn Albert, PhD), P30 AG062421-01 (PI Bradley Hyman, MD, PhD), P30 AG062422-01 (PI Ronald Petersen, MD, PhD), P50 AG005138 (PI Mary Sano, PhD), P30 AG008051 (PI Thomas Wisniewski, MD), P30 AG013854 (PI Robert Vassar, PhD), P30 AG008017 (PI Jeffrey Kaye, MD), P30 AG010161 (PI David Bennett, MD), P50 AG047366 (PI Victor Henderson, MD, MS), P30 AG010129 (PI Charles DeCarli, MD), P50 AG016573 (PI Frank LaFerla, PhD), P30 AG062429-01(PI James Brewer, MD, PhD), P50 AG023501 (PI Bruce Miller, MD), P30 AG035982 (PI Russell Swerdlow, MD), P30 AG028383 (PI Linda Van Eldik, PhD), P30 AG053760 (PI Henry Paulson, MD, PhD), P30 AG010124 (PI John Trojanowski, MD, PhD), P50 AG005133 (PI Oscar Lopez, MD), P50 AG005142 (PI Helena Chui, MD), P30 AG012300 (PI Roger Rosenberg, MD), P30 AG049638 (PI Suzanne Craft, PhD), P50 AG005136 (PI Thomas Grabowski, MD), P30 AG062715-01 (PI Sanjay Asthana, MD, FRCP), P50 AG005681 (PI John Morris, MD), P50 AG047270 (PI Stephen Strittmatter, MD, PhD).

## Author contributions

N.R.J., A.W., C.C., S.S., H.J.C., and H.P. designed the research. N.R.J., A.W., C.C., S.S., E.L., L.V., N.M., and C.A. performed the experiments. N.R.J., S.S., and H.P. analyzed the data. N.R.J., H.J.C., and H.P. wrote the manuscript. All authors edited the manuscript.

## Data and materials availability

All data generated or analyzed during this study involving the drug screen are included in this published article and its supplementary materials. The NACC data supporting the findings of this study are available on request from NACC at www.naccdata.org. All code generated during this study is included in the supplementary materials.

## Supplementary Materials

**Fig. S1.**
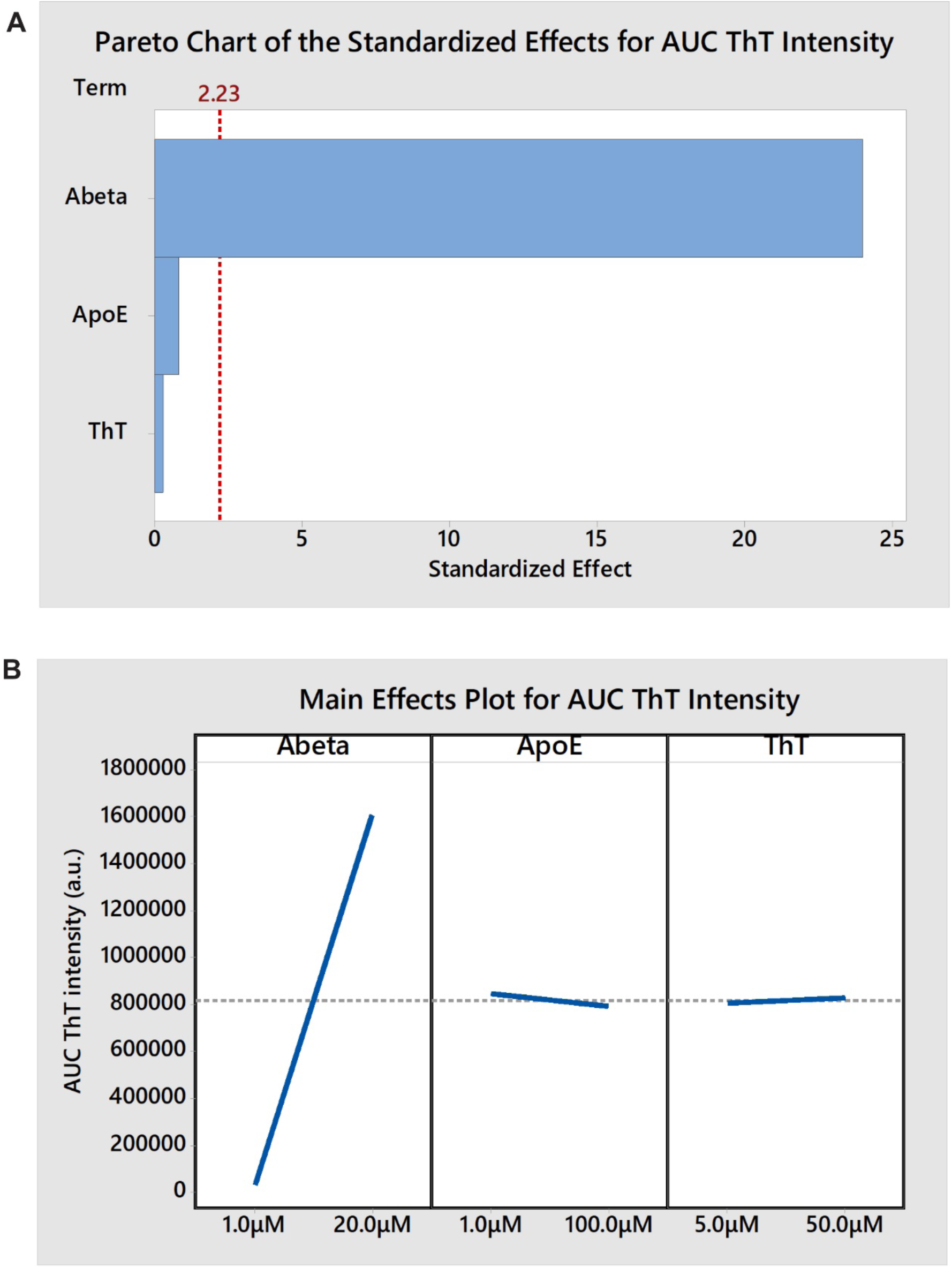
Half-fraction factorial design. Three reactant concentrations, Aβ, apoE4, and ThT, were varied in a half-fraction factorial design for a total of 2^3^/2 = 4 experimental conditions and one center point. Three technical replicates (wells) were tested per experimental condition. Experimental conditions and data are provided in Data file S1. (**A**) Pareto chart of the standardized effect for each reactant on the integrated area under the curve (AUC) of ThT intensity. The critical effect size for statistical significance (α = 0.05) is also shown at an effect size of 2.23 (red line). Aβ concentration had a large effect while the effects of apoE4 and ThT concentrations were insignificant. The interaction effects are confounded with the main effects and are therefore not shown. (**B**) Main effects plot showing the size and direction of each effect on the AUC of ThT intensity. As Aβ concentration increased from 1 µM to 20 µM the AUC of ThT intensity increased from 0 to approximately 1.6 x 10^6^ a.u., while apoE4 and ThT concentrations had no significant effects.

**Fig. S2.**
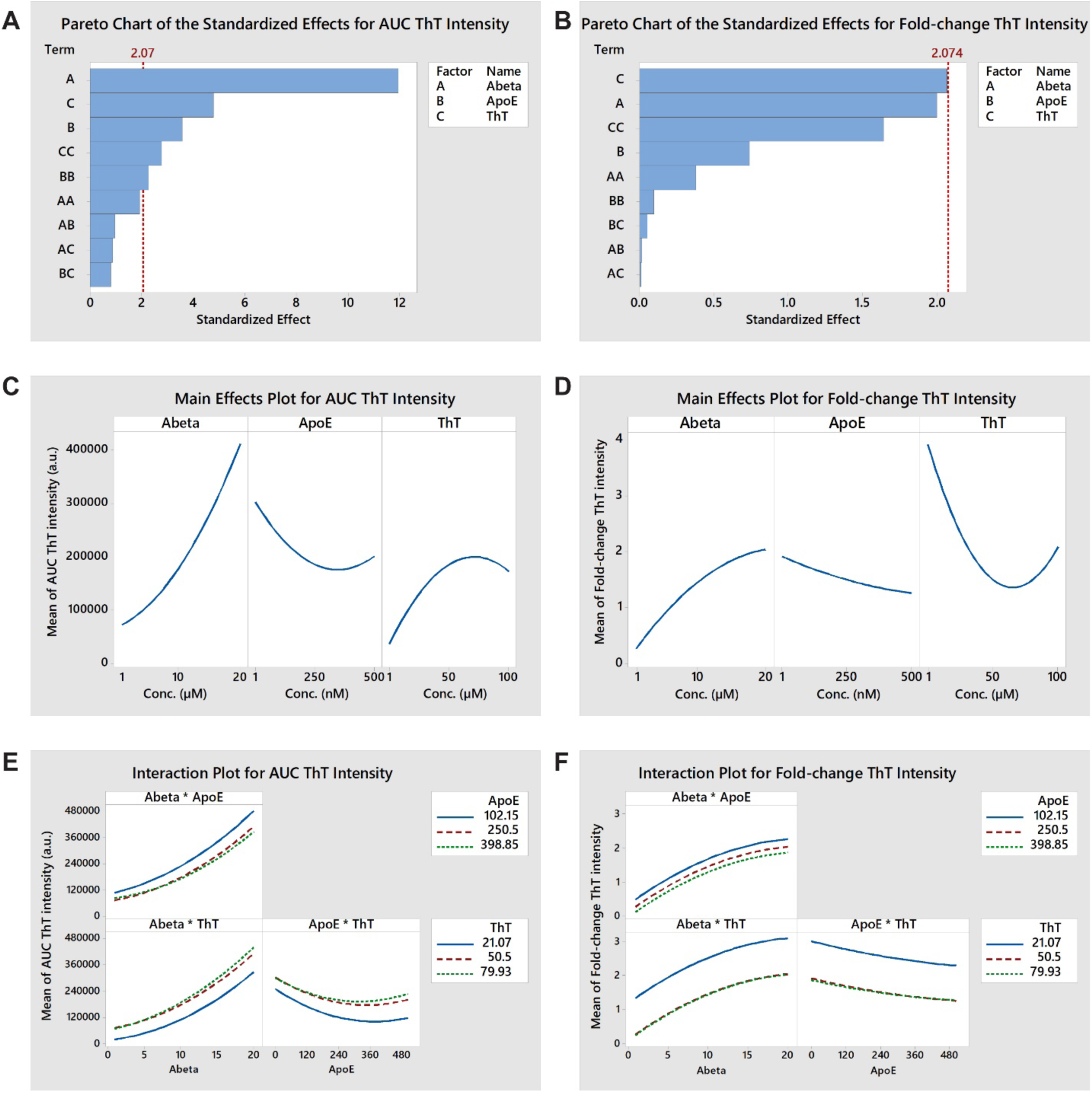
Central composite response surface design #1. Three reactant concentrations, Aβ, apoE4, and ThT, were varied in a central composite design using 2^3^ = 8 corner points, 2*3 = 6 axial points, and one center point. An optimized design space was determined based on the results of the previous factorial experiment. Two replicates (wells) were tested per experimental condition and four replicates of the center point. Experimental conditions and data are provided in Data file S1. (**A, B**) Pareto charts showing the standardized effect for the main (A, B, C), quadratic (AA, BB, CC), and interaction effects (AB, BC, AC) on the AUC and the fold-change of ThT intensity, respectively. The critical effect size for statistical significance (α = 0.05) is also shown at an effect size of 2.074 (red line). All three variables had large main and quadratic effects on the AUC of ThT intensity, while all interaction effects were negligible. ThT and Aβ concentrations had large main and quadratic effects on the fold-change in ThT intensity, while the effects of apoE concentration and all interaction effects were much smaller. (**C, D**) Main effects plots showing the combination of main and quadratic effects of each reactant on the AUC and the fold-change of ThT intensity, respectively. High Aβ concentration, low apoE4 concentration, and an intermediate ThT concentration maximized both the AUC, and the fold-change, of ThT intensity. (**E, F**) Interaction plots showing the interaction effect of each reactant pair on the AUC and the fold-change of ThT intensity, respectively. No significant interactions were observed, which is evidenced by the similar shapes of the response curves for all reactant concentrations in each interaction plot.

**Fig. S3.**
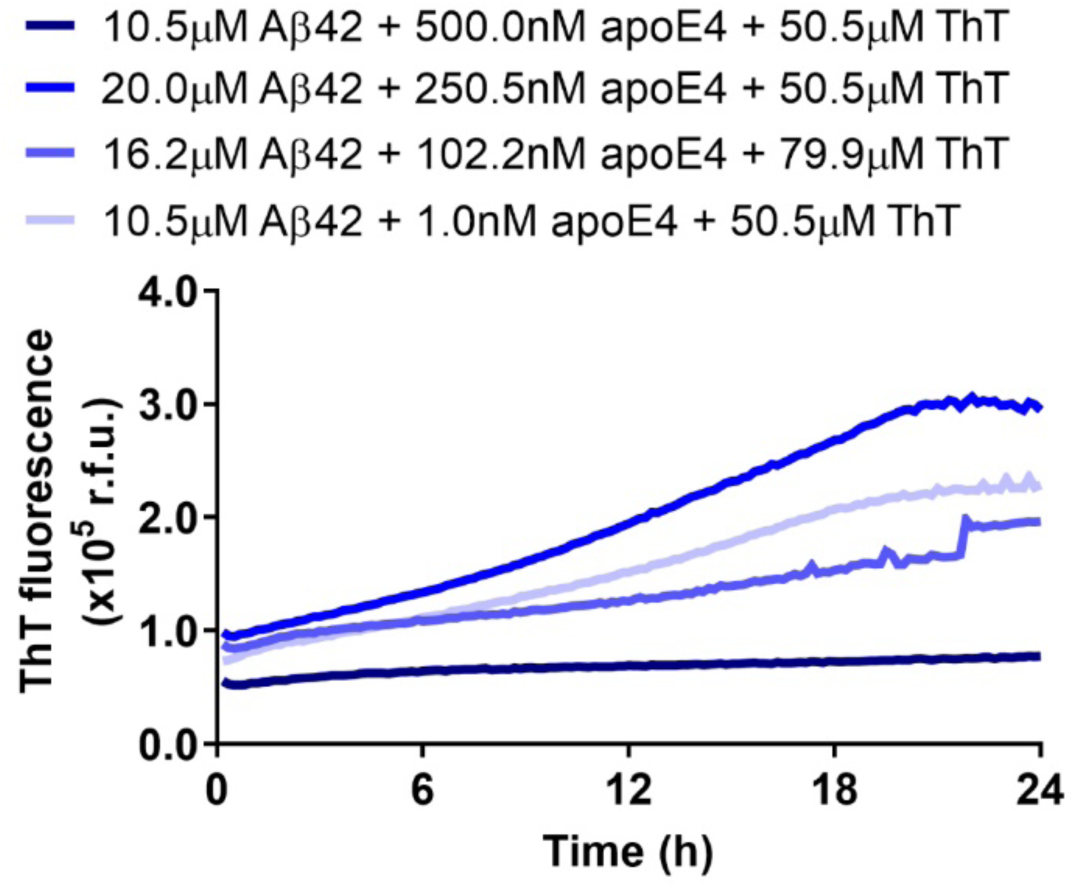
Effect of apoE4 concentration on Aβ42 fibrillization. Concentrations of Aβ42, apoE4, and ThT were varied in a response surface design. The fibrillization assay was run in a 384-well plate and was analyzed for ThT fluorescence over a 24 h period. Several groups were plotted to demonstrate the effects of the different concentrations of apoE4 on ThT fluorescence over time. The complete results are provided in Data file S1. The data represent the mean of n = 3−4 wells per group.

**Fig. S4.**
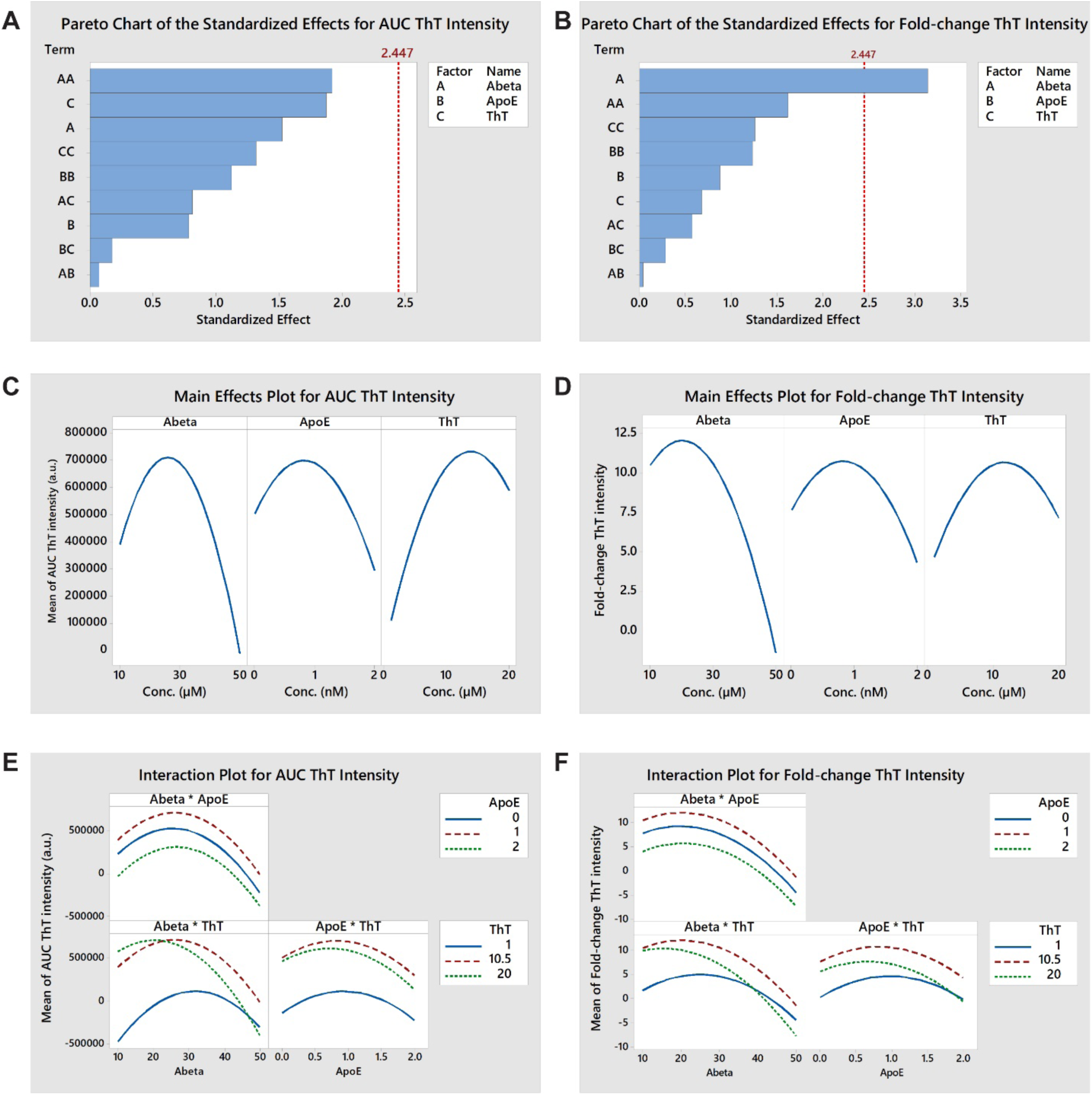
Central composite response surface design #2. Three reactant concentrations, Aβ, apoE4, and ThT, were varied in a central composite design using 2^3^ = 8 corner points, 2*3 = 6 axial points, and one center point. An optimized design space was determined based on the results of the previous response surface experiment. Three replicates (wells) were tested per experimental condition and six replicates of the center point, and the entire experiment was repeated in two independent experiments (blocks). Experimental conditions and data are provided in Data file S1. (**A, B**) Pareto charts showing the standardized effect for the main (A, B, C), quadratic (AA, BB, CC), and interaction effects (AB, BC, AC) on the AUC and the fold-change of ThT intensity, respectively. The critical effect size for statistical significance (α = 0.05) is also shown at an effect size of 2.447 (red line). Aβ and ThT had large main and quadratic effects on the AUC of ThT intensity, while the effect of apoE was smaller. Aβ had the largest main and quadratic effects on the fold-change in ThT intensity, while the effects of apoE and ThT were smaller. (**C, D**) Main effects plots showing the combination of main and quadratic effects of each reactant on the AUC and the fold-change of ThT intensity, respectively. Intermediate concentrations of Aβ, apoE4, and ThT maximized both the AUC and the fold-change of ThT intensity. (**E, F**) Interaction plots showing the interaction effect of each reactant pair on the AUC and the fold-change of ThT intensity, respectively. An interaction between Aβ and ThT concentrations (AC) was observed to have a moderate effect on both the AUC and the fold-change of ThT intensity, which is evidenced by the response curves for different reactant concentrations crossing one another. This moderate effect caused both responses to peak at lower Aβ concentrations when the ThT concentration was 20 μM compared to 10.5 μM. However, the interaction effect did not change the conclusions about the dominant main and quadratic effects of Aβ and ThT seen in the main effects plot.

**Fig. S5.**
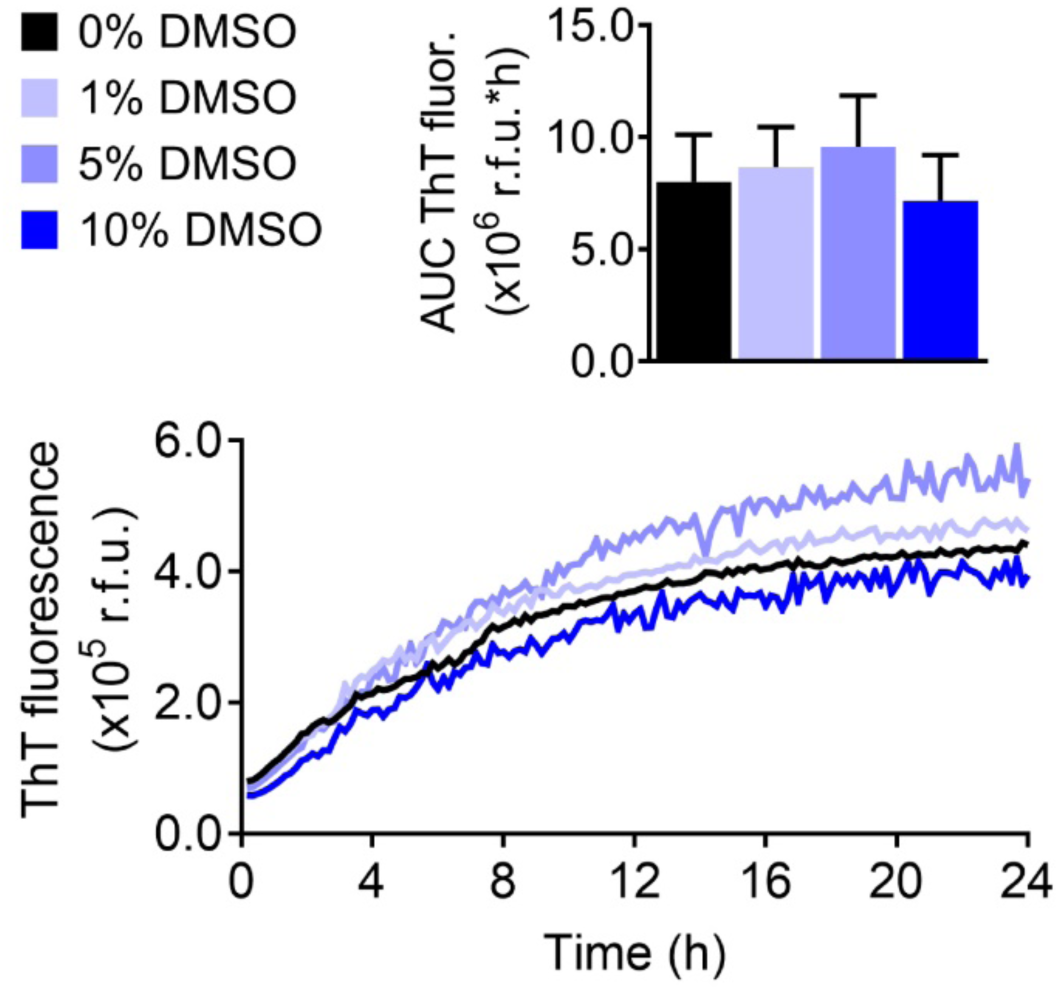
Effect of DMSO in the optimized apoE4-Aβ fibrillization assay. The effects of DMSO at 0, 1, 5, and 10% (v/v) on apoE4-catalyzed Aβ42 fibrillization were evaluated. The data represent the mean ± SD of n = 8 wells per group.

**Fig. S6.**
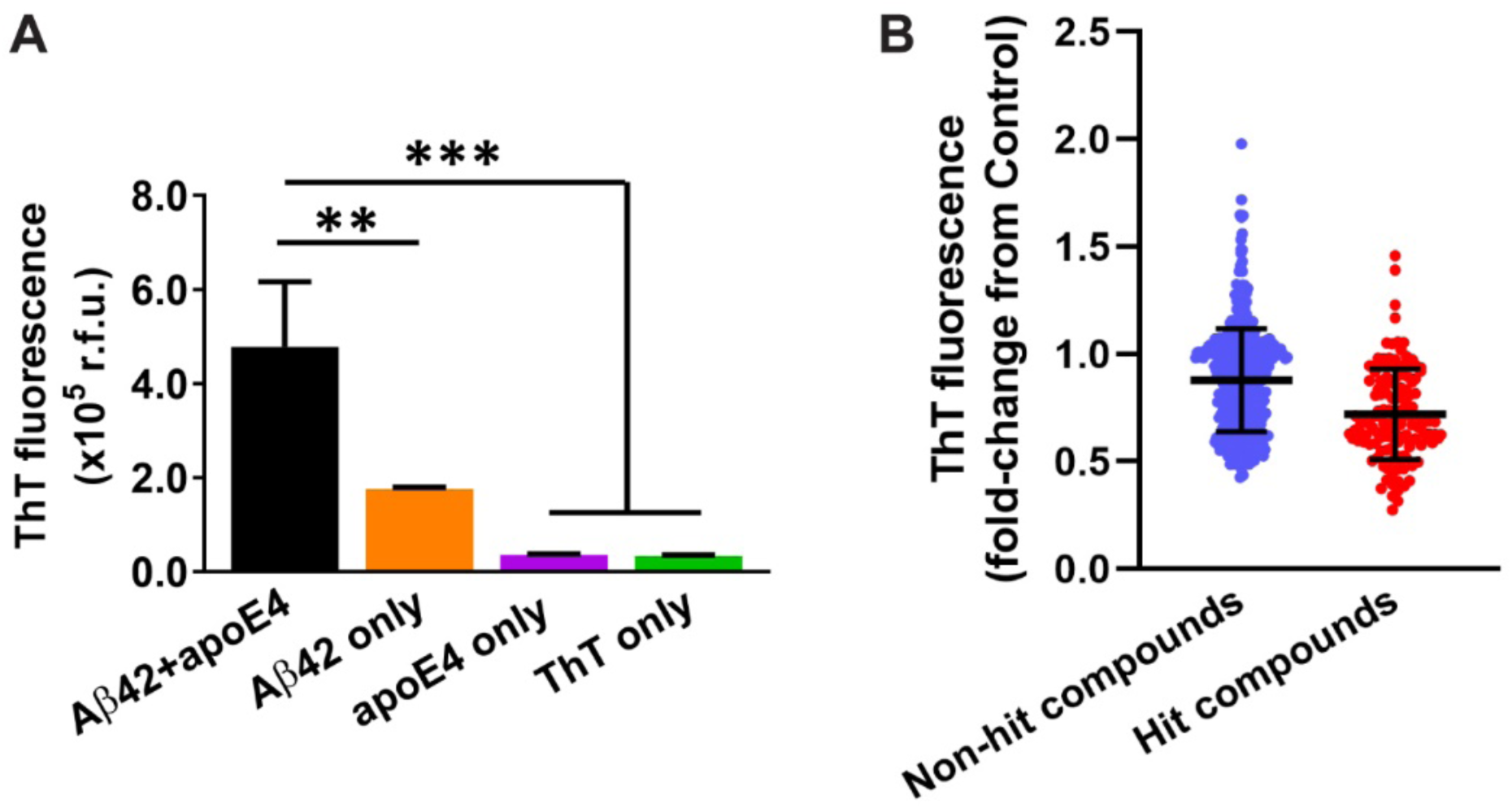
Exploratory drug screen. The apoE4/Aβ42 fibrillization assay was performed in an endpoint fashion in the exploratory screen. To set up the fibrillization assay, Aβ42 (2 μM) and apoE4 (20 nM) were combined in water in a 96-well plate and incubated for 15 mins. ThT and glycine were added and incubated for 10 mins, and then fluorescence was measured at λ_ex_ = 440 nm, λ_em_ = 490 nm. (**A**) Under these conditions, Aβ42+apoE4 resulted in significantly greater ThT fluorescence than Aβ42, apoE4, or ThT alone. The data represent the mean ± SD of n = 3 wells per group. Statistical significance is indicated as \*\**P* < 0.01, \*\*\**P* < 0.001 by one-way ANOVA. (**B**) In the exploratory screen, compounds (2 μM), or DMSO as the control, were initially incubated with Aβ42 and apoE4, and the ThT intensity for each compound was normalized to the control group on the same plate. A total of 595 compounds from the NCC library were evaluated in the exploratory screen, with 134 being identified as hit compounds (red dots) and 461 being identified as non-hit compounds (blue dots). Each data point represents a single compound tested in n = 3 wells and averaged, and the black lines indicate the mean ± SD for each group.

**Fig. S7.**
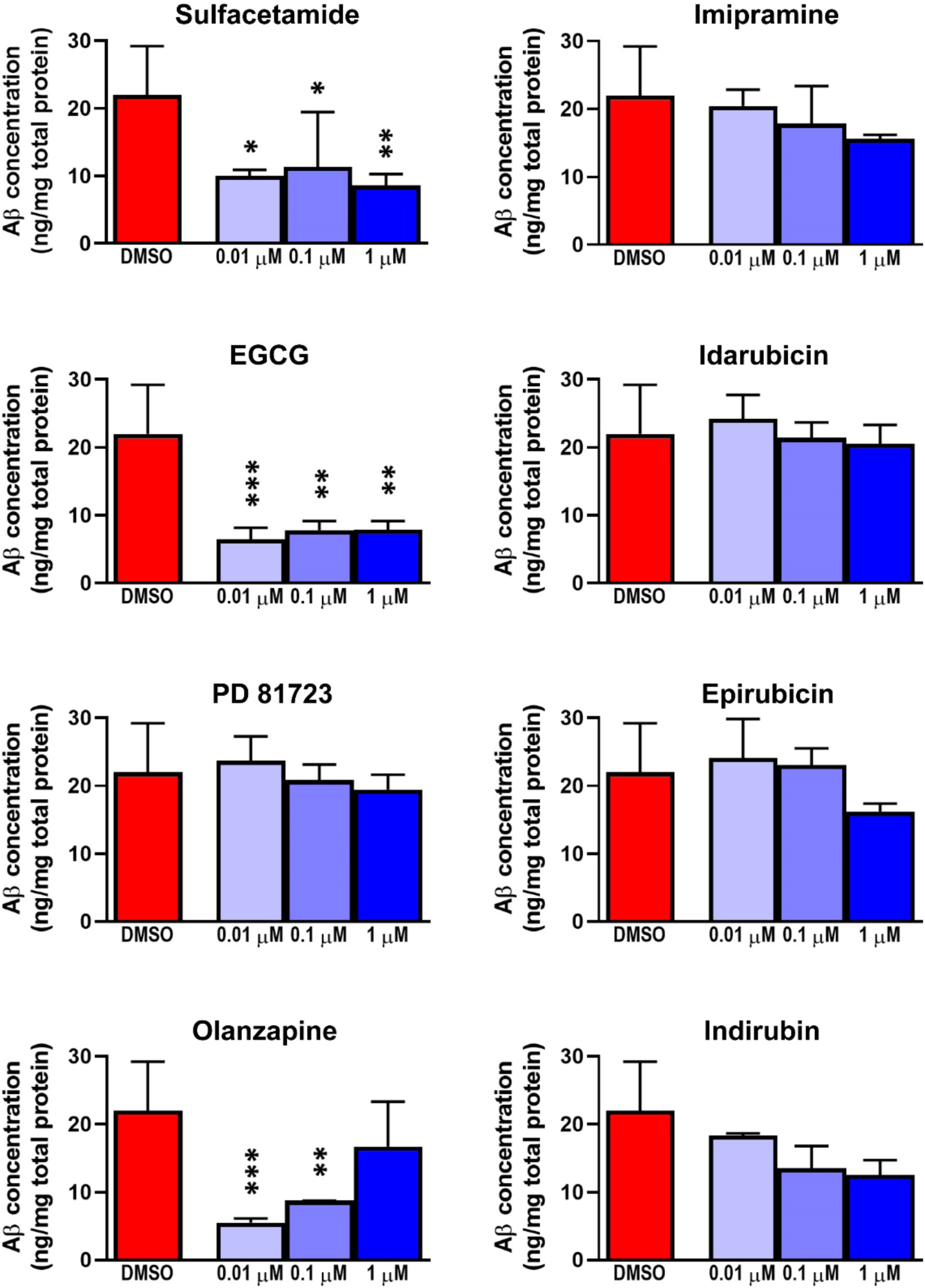
Aβ levels in conditioned medium of 5xFAD mouse neurons. Aβ42 concentrations were measured in the conditioned medium of 5xFAD mouse neurons at 9 dpe to each hit compound by enzyme-linked immunosorbent assay (ELISA). The data represent the mean ± SD of n = 6 wells for the DMSO control and n = 3 wells per concentration for compounds. Statistical significance is indicated as \**P* < 0.05, \*\**P* < 0.01, and \*\*\**P* < 0.001 compared to the DMSO control by one-way ANOVA.

**Fig. S8.**
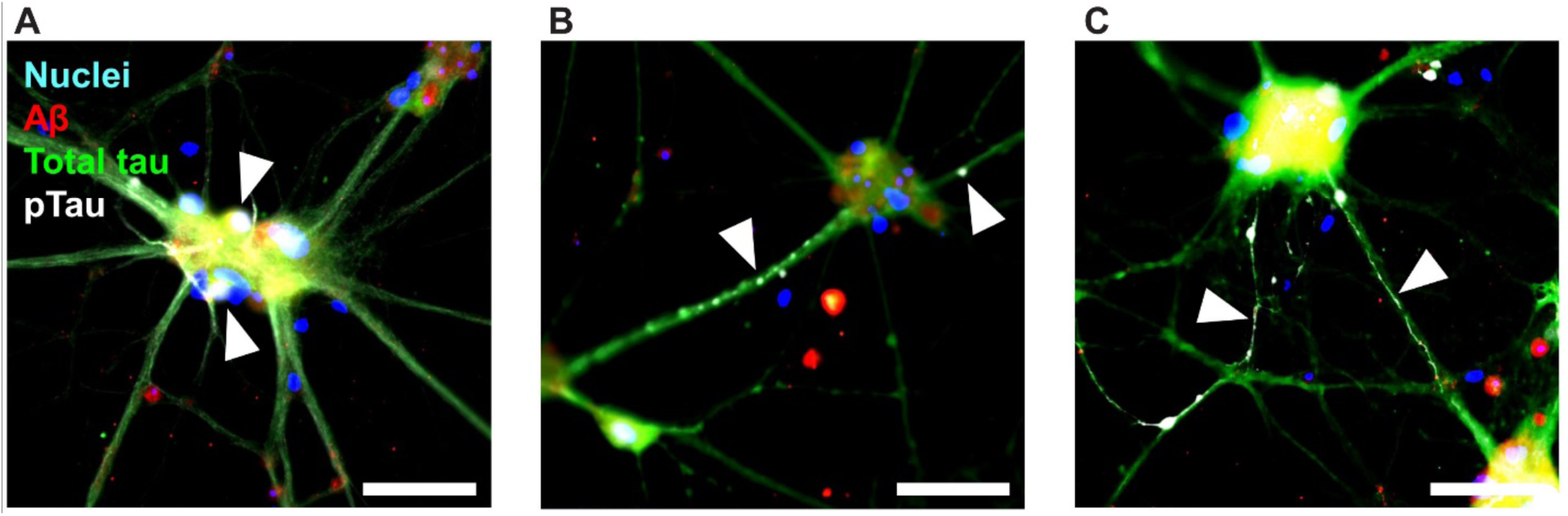
pTau neuropathological features observed in TgF344-AD primary rat neurons. Representative ICC images of neurons at 14 dpe to apoE4 and Aβ42, treated with DMSO only as a control, and labeled for Aβ (red), total tau (green), pTau [S202/T205] (white), and cell nuclei (blue). Characteristic pTau neuropathological features were observed including (**A**) intracellular and extracellular puncta, (**B**) axonal blebbing, and (**C**) neuropil thread-like structures. Arrowheads indicate respective pTau neuropathological features. Scale bars = 50 μm.

**Fig. S9.**
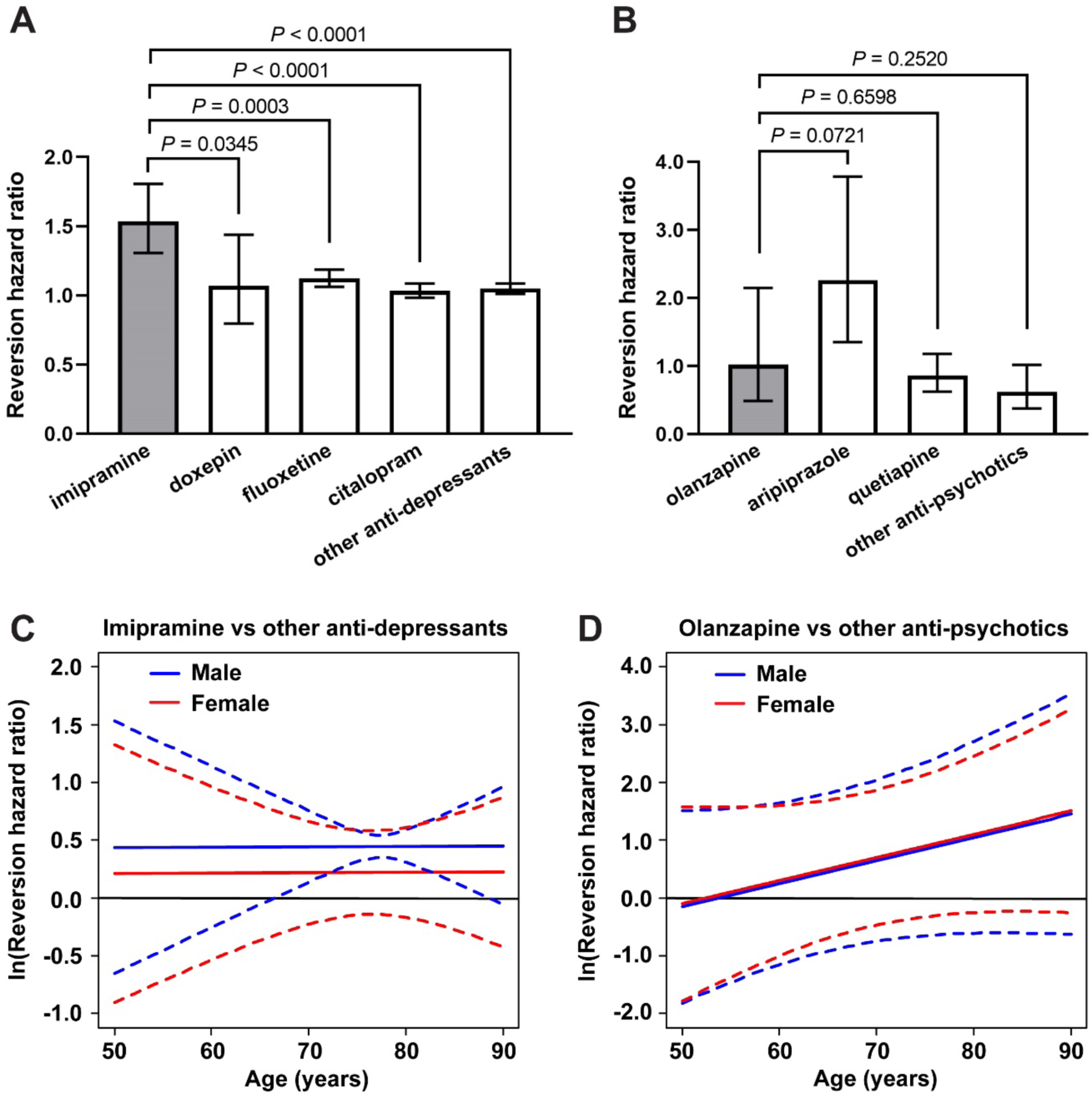
Mixed medication and interaction models evaluating clinical diagnosis reversion. (**A**) The hazard ratio of clinical diagnosis reversion toward normal was plotted comparing the cumulative drug exposure of imipramine, doxepin, fluoxetine, citalopram, or all other anti-depressants to being off the medication, in the same subjects. The data indicate the hazard ratio ± 95% CI. Imipramine was compared to each other group and all *P* values are shown. (**B**) The hazard ratio of clinical diagnosis reversion toward normal was plotted comparing the effect of being on olanzapine, aripiprazole, quetiapine, or all other anti-psychotics to being off the medication, in the same subjects. The data indicate the hazard ratio ± 95% CI. Olanzapine was compared to each other group and all *P* values are shown. (**C**) Imipramine was compared to other anti-depressant medications for the potential effect of cumulative drug exposure on the hazard ratio of clinical diagnosis reversion toward normal, with age and sex considered as interaction variables. The data represent the natural log of the HR (solid lines) and 95% CI (dotted lines). Statistical significance is reached when the 95% CI does not include zero, which occurs from 66.5−88.5 years of age in males. (**D**) Olanzapine was compared to other anti-psychotic medications for the potential effect of being on the medication on the hazard ratio of clinical diagnosis reversion, with age and sex considered as interaction variables. The data represent the natural log of the HR (solid lines) and 95% CI (dotted lines).

**Table S1.**
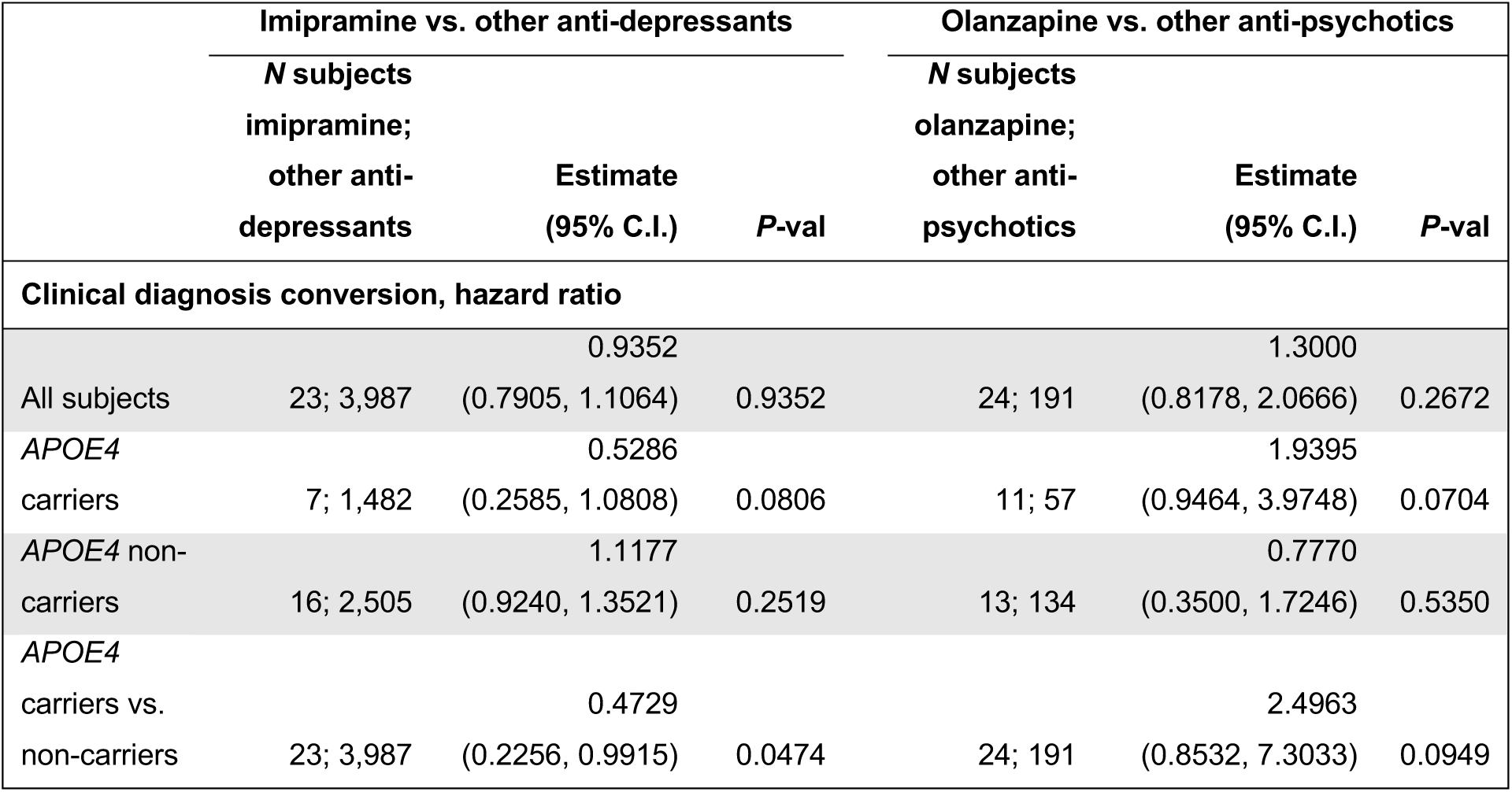
Retrospective analysis of NACC dataset for clinical diagnosis conversion. The cumulative exposure of imipramine and other anti-depressants were compared, and the on/off status of olanzapine and other anti-psychotics were compared, using Cox proportional hazard ratio analysis for statistical comparisons. Only subjects who reported use of a medication prior to clinical diagnosis conversion were included. Test statistics and degrees of freedom are provided in Data file S3.

**Data file S1. Design of experiments (DOE) assay optimization data.** Experimental conditions and raw data for (**A**) Half-fraction factorial experiment, (**B**) Response surface experiment #1, and (**C**) Response surface experiment #2. The RunOrder column indicates randomized order in which the different experimental conditions were prepared in the wells of a plate. The CenterPt column indicates whether the experimental condition is a center point (0) or not (1). The PtType column indicates whether the experimental condition is a corner or axial point (−1 or 1), or a center point (0). The Blocks column indicates whether the experimental condition was included in a single plate run on one day (1), or was included in a second plate repeating the entire experiment on a different day (2). The AUC ThT intensity column indicates the integrated area under the curve of ThT fluorescence intensity measured by the plate reader in arbitrary units (a.u.) over the entire experiment duration. The Fold-change ThT intensity column indicates the fold-change in ThT fluorescence intensity from the beginning to the end of the experiment.

**Data file S2. Exploratory drug screen results.** (**A**) Detailed information about the 134 hit compounds identified in the exploratory screen. (**B**) Blood-brain barrier (BBB) permeability for the 134 hit compounds identified in the exploratory screen, as determined by a literature search. Positive BBB qualities were found for 87 compounds (labeled in green), while negative BBB qualities (labeled in red) or no information (labeled in yellow) were found for 41 compounds and 6 compounds, respectively.

**Data file S3. Summary statistics of NACC data analyses.** (**A**, **B**) Subject information for MMSE models comparing (**A**) imipramine and other anti-depressants or (**B**) olanzapine and other anti-psychotics, including age, sex, baseline MMSE score, and drug exposure time. (**C**, **D**) Subject information for clinical diagnosis reversion models comparing (**C**) imipramine and other anti-depressants or (**D**) olanzapine and other anti-psychotics, including age, sex, baseline MMSE score, drug exposure time, number of subjects with reversions, and number of reversions per subject. (**E**, **F**) Subject information for clinical diagnosis conversion models comparing (**E**) imipramine and other anti-depressants or (**F**) olanzapine and other anti-psychotics, including age, sex, baseline MMSE score, drug exposure time, number of subjects with conversions, and number of conversions per subject. (**G**, **H**) Subject information for multiple medications models comparing (**G**) imipramine, doxepin, fluoxetine, citalopram, and all other anti-depressants or (**H**) olanzapine, aripiprazole, quetiapine, and all other anti-psychotics, including age, sex, baseline MMSE score, drug exposure time, number of subjects with reversions, and number of reversions per subject. (**I**) Complete test statistics and degrees of freedom for all statistical tests.

**Data file S4. Custom computer code generated for NACC data analysis.** Computer code written in SAS and used to perform all statistical analyses of NACC data is provided in a text file. Computer code written in R and used to generate plots of NACC data is provided as an R file.

