## Supplementary material for "Imipramine and olanzapine block apoE4-catalyzed polymerization of Aβ and show evidence of improving Alzheimer’s disease cognition"

**Supplementary Materials**

A

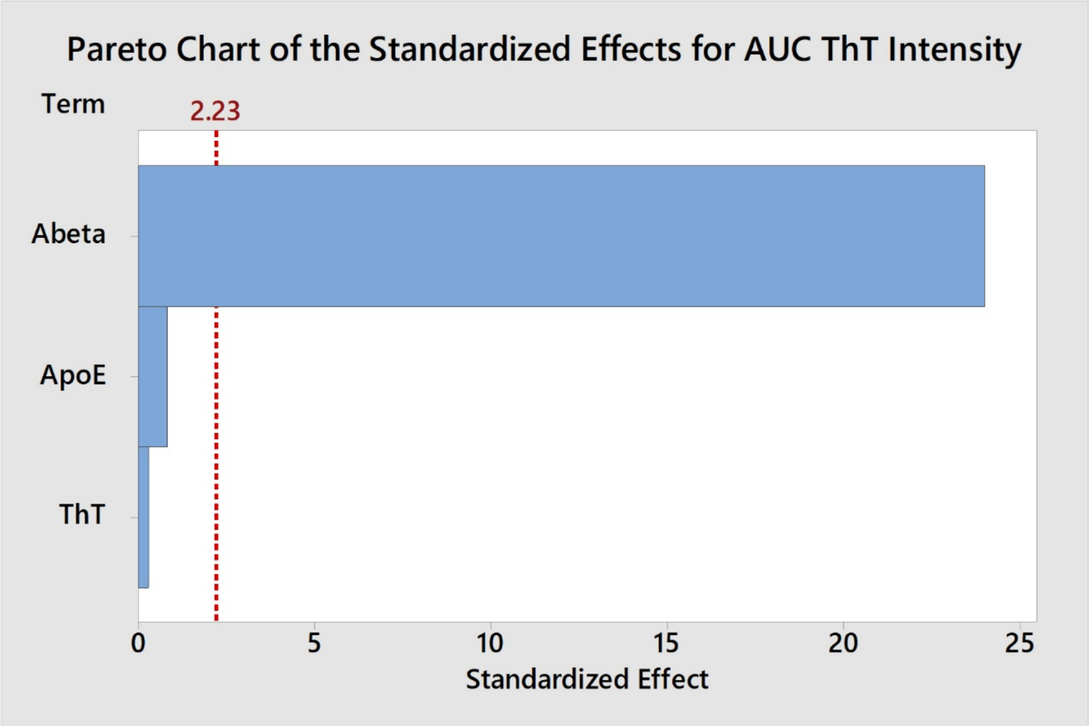


B

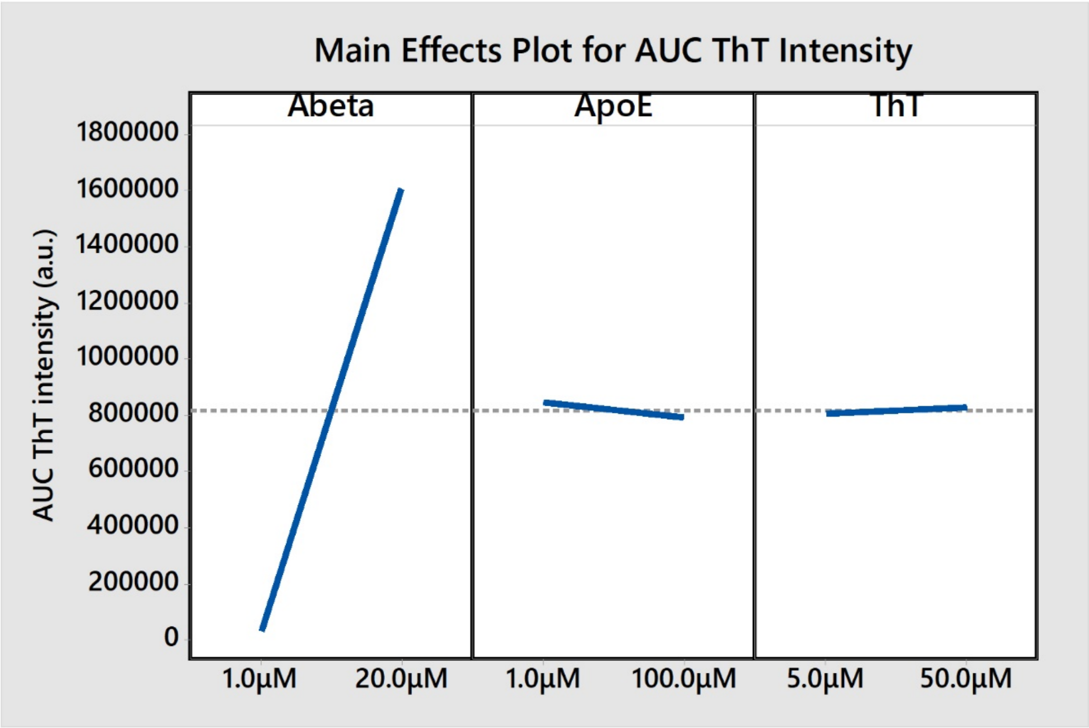


**Fig. S1. Half-fraction factorial design.** Three reactant concentrations, Aβ, apoE4, and ThT, were varied in a half-fraction factorial design for a total of $2^3/2 = 4$ experimental conditions and one center point. Three technical replicates (wells) were tested per experimental condition. Experimental conditions and data are provided in Data file S1. (**A**) Pareto chart of the standardized effect for each reactant on the integrated area under the curve (AUC) of ThT intensity. The critical effect size for statistical significance ($\alpha = 0.05$) is also shown at an effect size of 2.23 (red line). Aβ concentration had a large effect while the effects of apoE4 and ThT concentrations were insignificant. The interaction effects are confounded with the main effects and are therefore not shown. (**B**) Main effects plot showing the size and direction of each effect on the AUC of ThT intensity. As Aβ concentration increased from 1 μM to 20 μM the AUC of ThT intensity increased from 0 to approximately $1.6 \times 10^6$ a.u., while apoE4 and ThT concentrations had no significant effects.

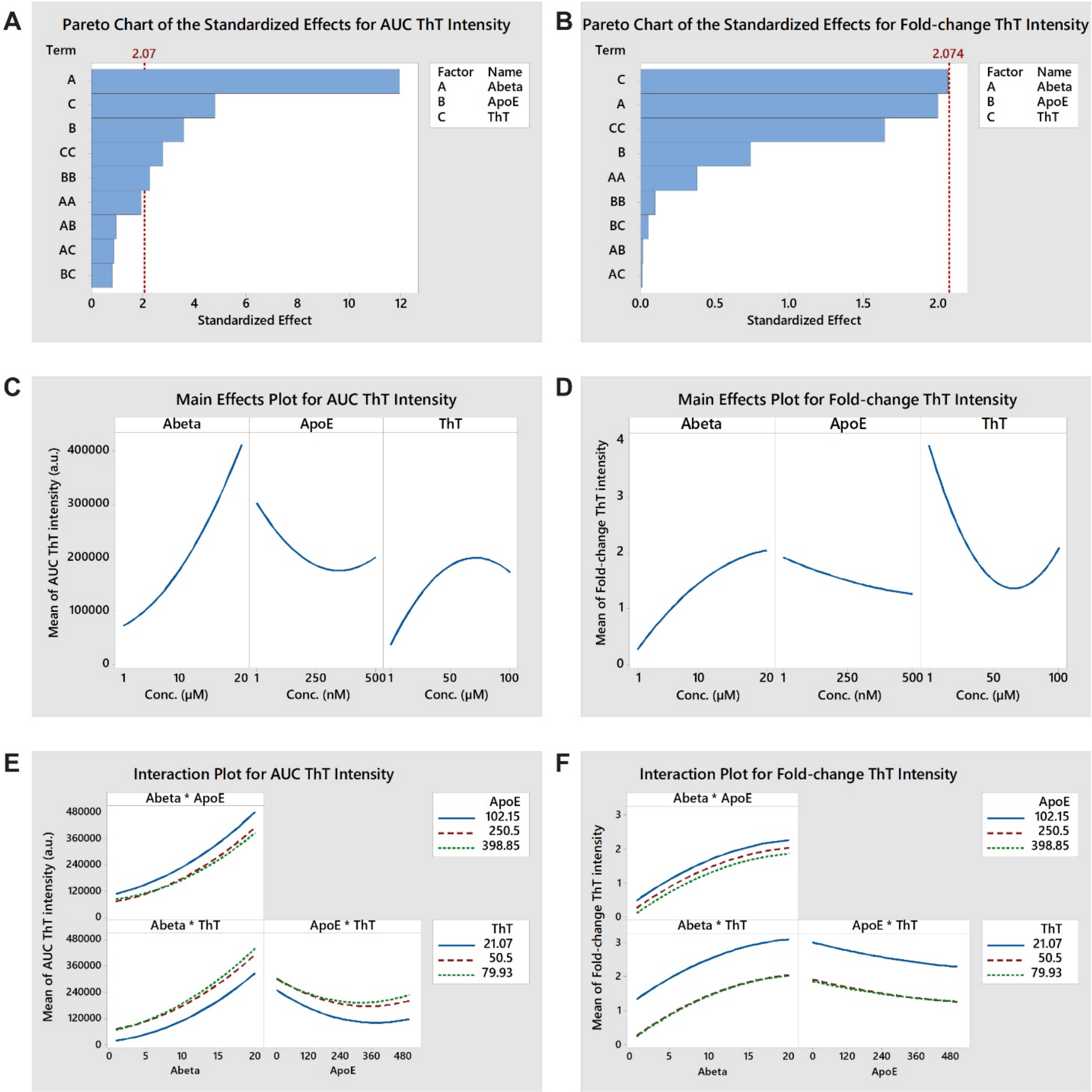


**Fig. S2. Central composite response surface design #1.** Three reactant concentrations, Aβ, apoE4, and ThT, were varied in a central composite design using $2^3 = 8$ corner points, $2*3 = 6$ axial points, and one center point. An optimized design space was determined based on the results of the previous factorial experiment. Two replicates (wells) were tested per experimental condition and four replicates of the center point. Experimental conditions and data are provided in Data file S1. (**A, B**) Pareto charts showing the standardized effect for the main (A, B, C), quadratic (AA, BB, CC), and interaction effects (AB, BC, AC) on the AUC and the fold-change of ThT intensity, respectively. The critical effect size for statistical significance ($\alpha = 0.05$) is also shown at an effect size of 2.074 (red line). All three variables had large main and quadratic effects on the AUC of ThT intensity, while all interaction effects were negligible. ThT and Aβ concentrations had large main and quadratic effects on the fold-change in ThT intensity, while the effects of apoE concentration and all interaction effects were much smaller. (**C, D**) Main effects plots showing the combination of main and quadratic effects of each reactant on the AUC and the fold-change of ThT intensity, respectively. High Aβ concentration, low apoE4 concentration, and an intermediate ThT concentration maximized both the AUC, and the fold-change, of ThT intensity. (**E, F**) Interaction plots showing the interaction effect of each reactant pair on the AUC and the fold-change of ThT intensity, respectively. No significant interactions were observed, which is evidenced by the similar shapes of the response curves for all reactant concentrations in each interaction plot.

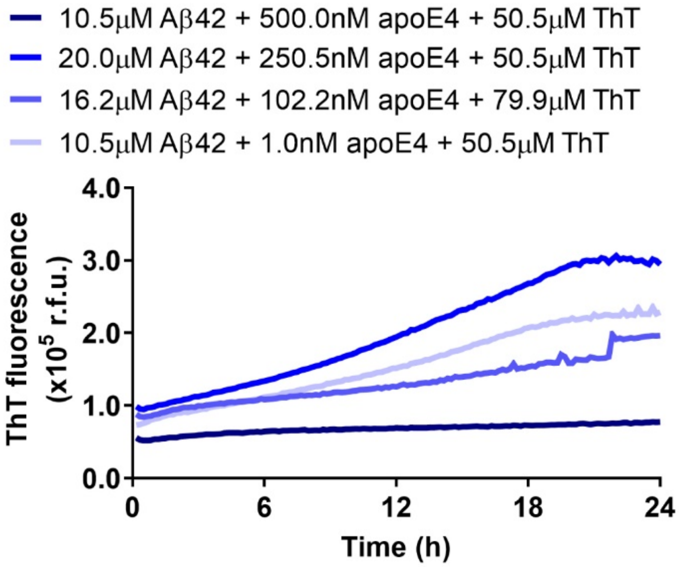


**Fig. S3. Effect of apoE4 concentration on Aβ42 fibrillization.** Concentrations of Aβ42, apoE4, and ThT were varied in a response surface design. The fibrillization assay was run in a 384-well plate and was analyzed for ThT fluorescence over a 24 h period. Several groups were plotted to demonstrate the effects of the different concentrations of apoE4 on ThT fluorescence over time. The complete results are provided in Data file S1. The data represent the mean of n = 3−4 wells per group.

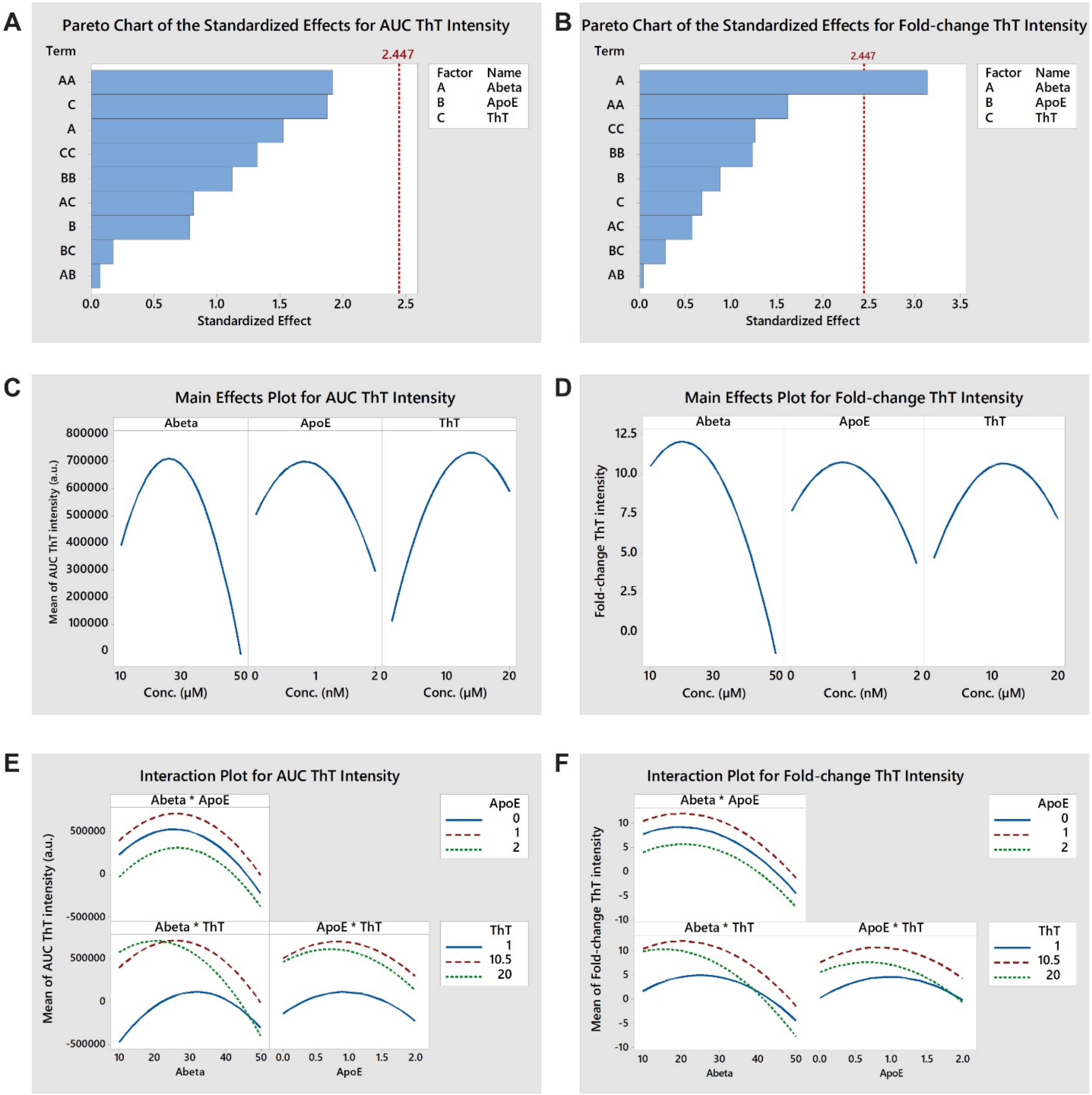


**Fig. S4. Central composite response surface design #2.** Three reactant concentrations, Aβ, apoE4, and ThT, were varied in a central composite design using $2^3 = 8$ corner points, $2*3 = 6$ axial points, and one center point. An optimized design space was determined based on the results of the previous response surface experiment. Three replicates (wells) were tested per experimental condition and six replicates of the center point, and the entire experiment was repeated in two independent experiments (blocks). Experimental conditions and data are provided in Data file S1. (**A, B**) Pareto charts showing the standardized effect for the main (A, B, C), quadratic (AA, BB, CC), and interaction effects (AB, BC, AC) on the AUC and the fold-change of ThT intensity, respectively. The critical effect size for statistical significance ($\alpha$ = 0.05) is also shown at an effect size of 2.447 (red line). Aβ and ThT had large main and quadratic effects on the AUC of ThT intensity, while the effect of apoE was smaller. Aβ had the largest main and quadratic effects on the fold-change in ThT intensity, while the effects of apoE and ThT were smaller. (**C, D**) Main effects plots showing the combination of main and quadratic effects of each reactant on the AUC and the fold-change of ThT intensity, respectively. Intermediate concentrations of Aβ, apoE4, and ThT maximized both the AUC and the fold-change of ThT intensity. (**E, F**) Interaction plots showing the interaction effect of each reactant pair on the AUC and the fold-change of ThT intensity, respectively. An interaction between Aβ and ThT concentrations (AC) was observed to have a moderate effect on both the AUC and the fold-change of ThT intensity, which is evidenced by the response curves for different reactant concentrations crossing one another. This moderate effect caused both responses to peak at lower Aβ concentrations when the ThT concentration was 20 μM compared to 10.5 μM. However, the interaction effect did not change the conclusions about the dominant main and quadratic effects of Aβ and ThT seen in the main effects plot.

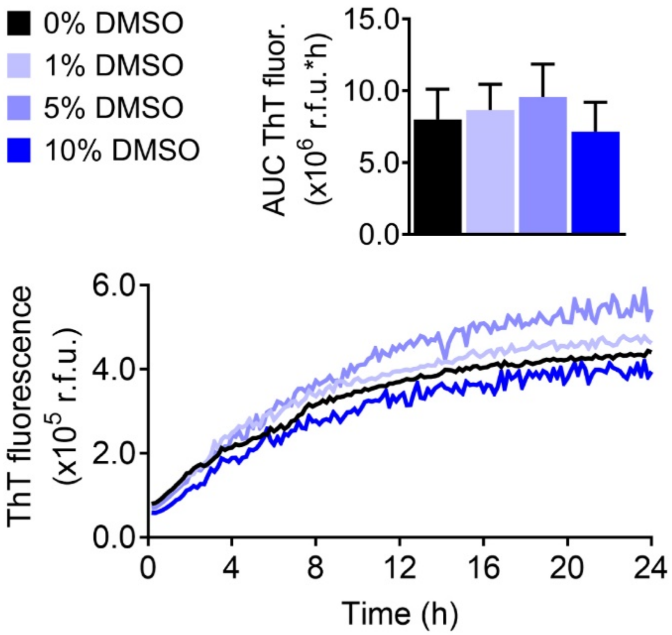


**Fig. S5. Effect of DMSO in the optimized apoE4-Aβ fibrillization assay.** The effects of DMSO at 0, 1, 5, and 10% (v/v) on apoE4-catalyzed Aβ42 fibrillization were evaluated. The data represent the mean ± SD of n = 8 wells per group.

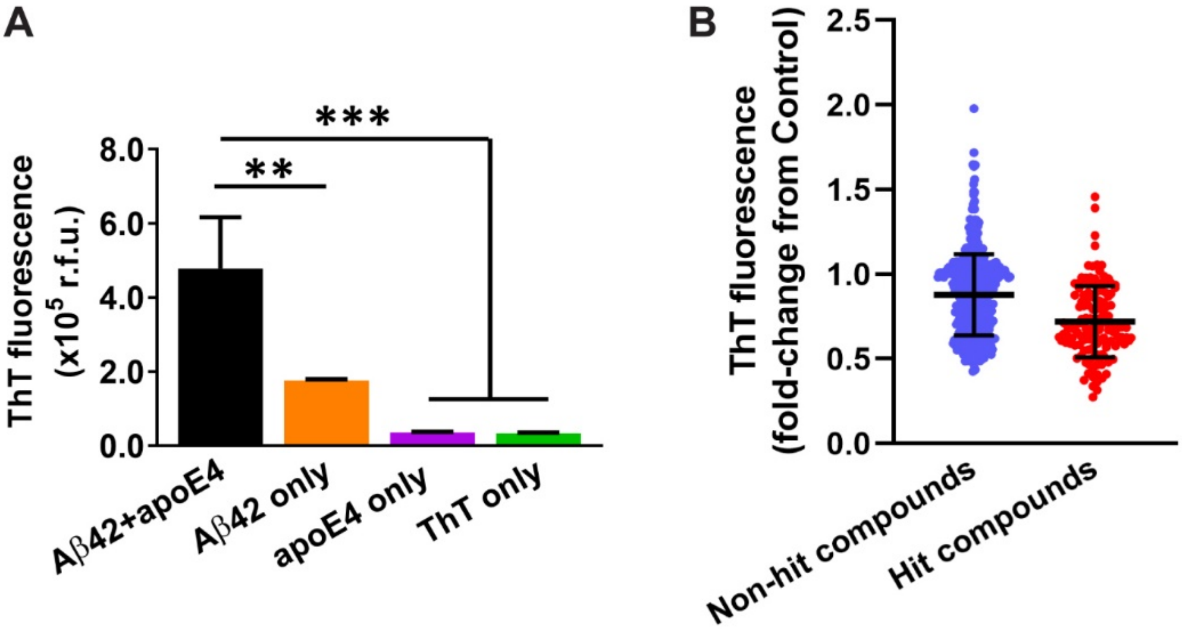


**Fig. S6. Exploratory drug screen.** The apoE4/Aβ42 fibrillization assay was performed in an endpoint fashion in the exploratory screen. To set up the fibrillization assay, Aβ42 (2 μM) and apoE4 (20 nM) were combined in water in a 96-well plate and incubated for 15 mins. ThT and glycine were added and incubated for 10 mins, and then fluorescence was measured at $\lambda_{ex}$ = 440 nm, $\lambda_{em}$ = 490 nm. (**A**) Under these conditions, Aβ42+apoE4 resulted in significantly greater ThT fluorescence than Aβ42, apoE4, or ThT alone. The data represent the mean ± SD of n = 3 wells per group. Statistical significance is indicated as $^{**}P < 0.01$, $^{***}P < 0.001$ by one-way ANOVA. (**B**) In the exploratory screen, compounds (2 μM), or DMSO as the control, were initially incubated with Aβ42 and apoE4, and the ThT intensity for each compound was normalized to the control group on the same plate. A total of 595 compounds from the NCC library were evaluated in the exploratory screen, with 134 being identified as hit compounds (red dots) and 461 being identified as non-hit compounds (blue dots). Each data point represents a single compound tested in n = 3 wells and averaged, and the black lines indicate the mean ± SD for each group.

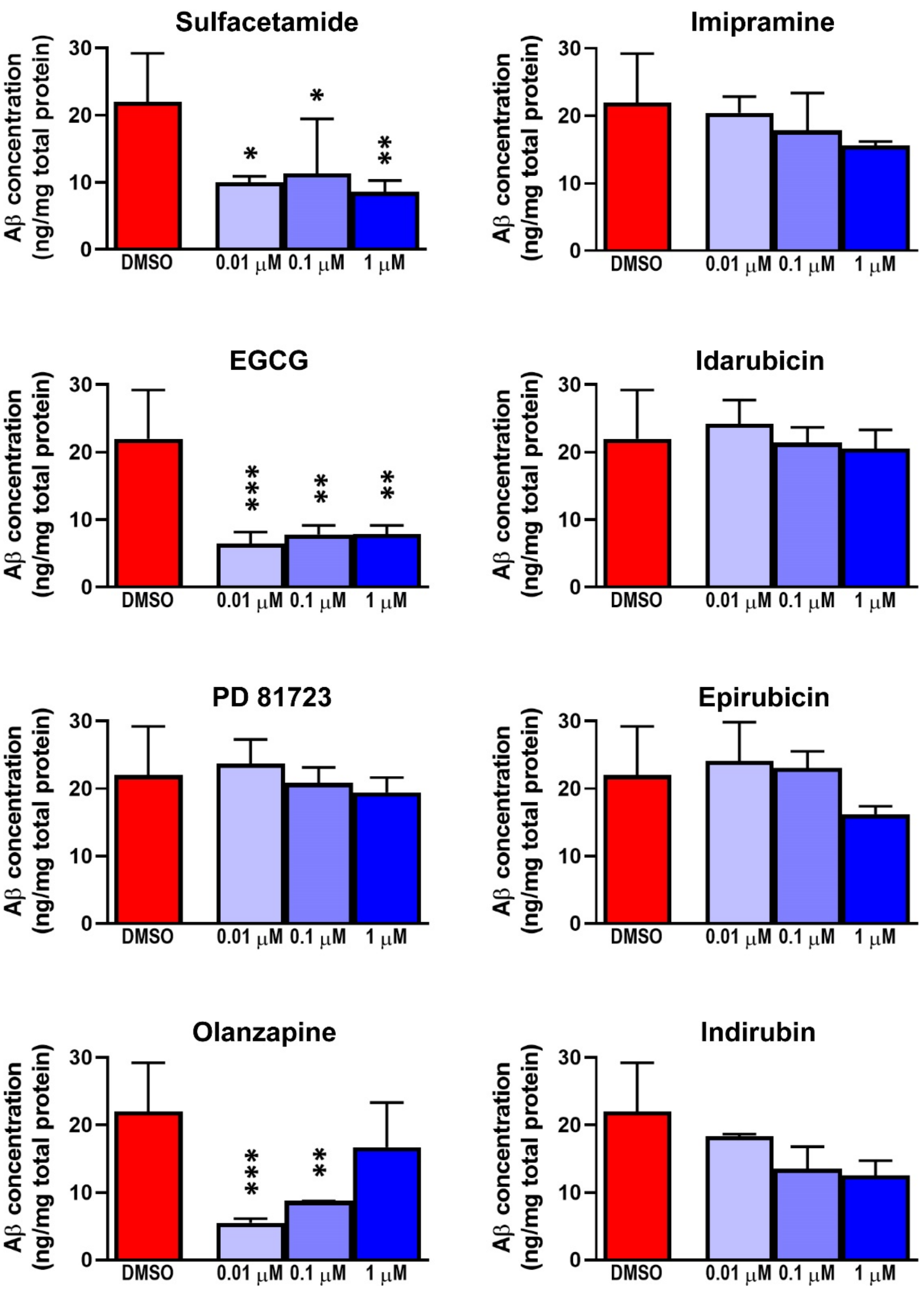


**Fig. S7. Aβ levels in conditioned medium of 5xFAD mouse neurons.** Aβ42 concentrations were measured in the conditioned medium of 5xFAD mouse neurons at 9 dpe to each hit compound by enzyme-linked immunosorbent assay (ELISA). The data represent the mean ± SD of n = 6 wells for the DMSO control and n = 3 wells per concentration for compounds. Statistical significance is indicated as $*P < 0.05$, $**P < 0.01$, and $***P < 0.001$ compared to the DMSO control by one-way ANOVA.

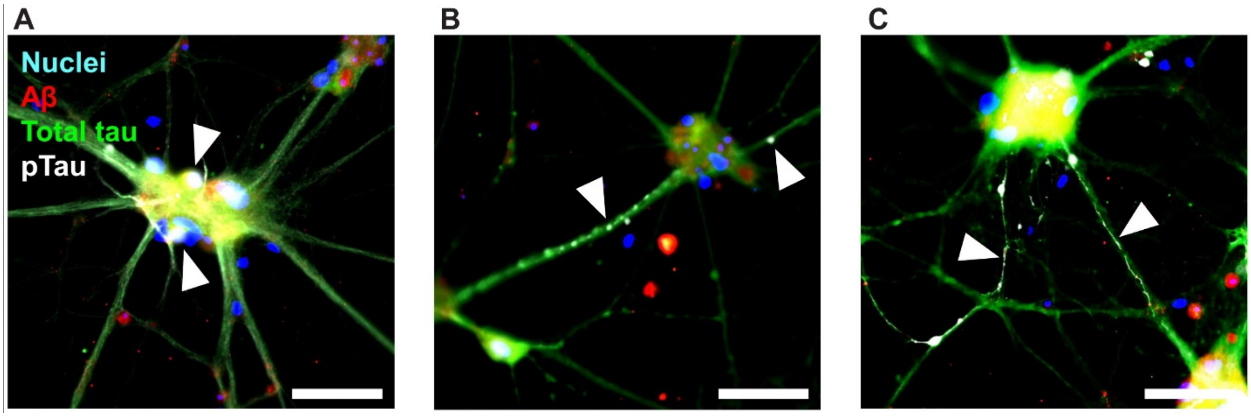


**Fig. S8. pTau neuropathological features observed in TgF344-AD primary rat neurons.** Representative ICC images of neurons at 14 dpe to apoE4 and Aβ42, treated with DMSO only as a control, and labeled for Aβ (red), total tau (green), pTau [S202/T205] (white), and cell nuclei (blue). Characteristic pTau neuropathological features were observed including (**A**) intracellular and extracellular puncta, (**B**) axonal blebbing, and (**C**) neuropil thread-like structures. Arrowheads indicate respective pTau neuropathological features. Scale bars = 50 μm.

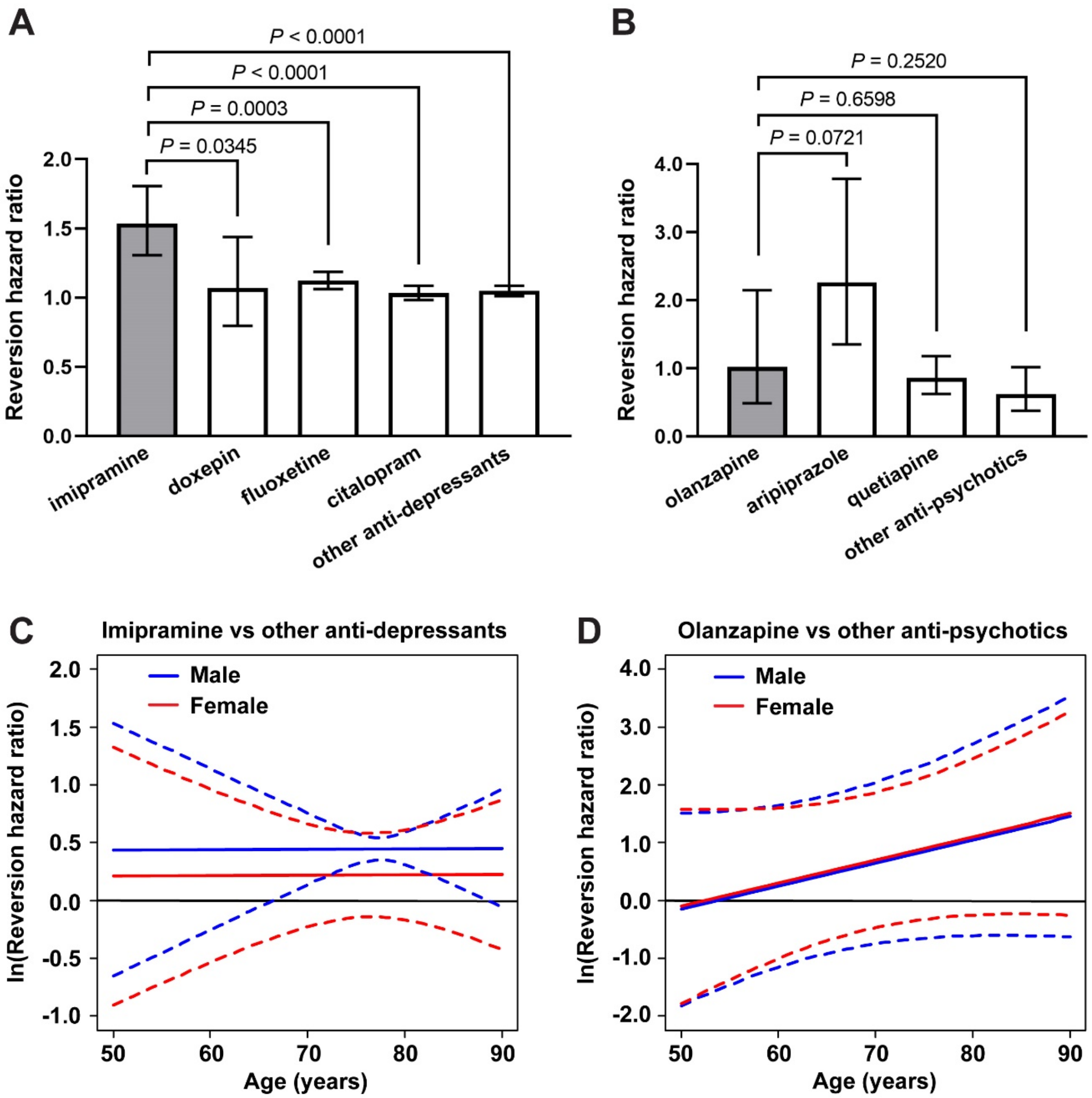


**Fig. S9. Mixed medication and interaction models evaluating clinical diagnosis reversion.** (**A**) The hazard ratio of clinical diagnosis reversion toward normal was plotted comparing the cumulative drug exposure of imipramine, doxepin, fluoxetine, citalopram, or all other anti-depressants to being off the medication, in the same subjects. The data indicate the hazard ratio ± 95% CI. Imipramine was compared to each other group and all *P* values are shown. (**B**) The hazard ratio of clinical diagnosis reversion toward normal was plotted comparing the effect of being on olanzapine, aripiprazole, quetiapine, or all other anti-psychotics to being off the medication, in the same subjects. The data indicate the hazard ratio ± 95% CI. Olanzapine was compared to each other group and all *P* values are shown. (**C**) Imipramine was compared to other anti-depressant medications for the potential effect of cumulative drug exposure on the hazard ratio of clinical diagnosis reversion toward normal, with age and sex considered as interaction variables. The data represent the natural log of the HR (solid lines) and 95% CI (dotted lines). Statistical significance is reached when the 95% CI does not include zero, which occurs from 66.5−88.5 years of age in males. (**D**) Olanzapine was compared to other anti-psychotic medications for the potential effect of being on the medication on the hazard ratio of clinical diagnosis reversion, with age and sex considered as interaction variables. The data represent the natural log of the HR (solid lines) and 95% CI (dotted lines).

| | Imipramine vs. other anti-depressants | | | Olanzapine vs. other anti-psychotics | | |
|---|---|---|---|---|---|---|
| | ***N* subjects imipramine; other anti-depressants** | **Estimate (95% C.I.)** | ***P*-val** | ***N* subjects olanzapine; other anti-psychotics** | **Estimate (95% C.I.)** | ***P*-val** |
| **Clinical diagnosis conversion, hazard ratio** | | | | | | |
| All subjects | 23; 3,987 | 0.9352 (0.7905, 1.1064) | 0.9352 | 24; 191 | 1.3000 (0.8178, 2.0666) | 0.2672 |
| *APOE4* carriers | 7; 1,482 | 0.5286 (0.2585, 1.0808) | 0.0806 | 11; 57 | 1.9395 (0.9464, 3.9748) | 0.0704 |
| *APOE4* non-carriers | 16; 2,505 | 1.1177 (0.9240, 1.3521) | 0.2519 | 13; 134 | 0.7770 (0.3500, 1.7246) | 0.5350 |
| *APOE4* carriers vs. non-carriers | 23; 3,987 | 0.4729 (0.2256, 0.9915) | 0.0474 | 24; 191 | 2.4963 (0.8532, 7.3033) | 0.0949 |

**Table S1. Retrospective analysis of NACC dataset for clinical diagnosis conversion.** The cumulative exposure of imipramine and other anti-depressants were compared, and the on/off status of olanzapine and other anti-psychotics were compared, using Cox proportional hazard ratio analysis for statistical comparisons. Only subjects who reported use of a medication prior to clinical diagnosis conversion were included. Test statistics and degrees of freedom are provided in Data file S3.
